# Precision of readout at the hunchback gene

**DOI:** 10.1101/063784

**Authors:** Jonathan Desponds, Huy Tran, Teresa Ferraro, Tanguy Lucas, Carmina Perez Romero, Aurelien Guillou, Cecile Fradin, Mathieu Coppey, Nathalie Dostatni, Aleksandra M. Walczak

## Abstract

The simultaneous expression of the *hunchback* gene in the multiple nuclei of the developing fly embryo gives us a unique opportunity to study how transcription is regulated in functional organisms. A recently developed MS2-MCP technique for imaging transcription in living *Drosophila* embryos allows us to quantify the dynamics of the developmental transcription process. The initial measurement of the morphogens by the *hunchback* promoter takes place during very short cell cycles, not only giving each nucleus little time for a precise readout, but also resulting in short time traces. Additionally, the relationship between the measured signal and the promoter state depends on the molecular design of the reporting probe. We develop an analysis approach based on tailor made autocorrelation functions that overcomes the short trace problems and quantifies the dynamics of transcription initiation. Based on life imaging data, we identify signatures of bursty transcription initiation from the *hunchback* promoter. We show that the precision of the expression of the *hunchback* gene to measure its position along the anterior-posterior axis is low both at the boundary and in the anterior even at cycle 13, suggesting additional post-translational averaging mechanisms to provide the precision observed in fixed material.

## I. INTRODUCTION

During development the different identities of cells are determined by sequentially expressing particular subsets of genes in different parts of the embryo. Proper development relies on the correct spatial-temporal assignment of cell types. In the fly embryo, the initial information about the position along the anterior-posterior (AP) axis is encoded in the exponentially decaying Bicoid gradient. The simultaneous expression of the Bicoid target gene *hunchback* in the multiple nuclei of the developing fly embryo gives us a unique opportunity to study how transcription is regulated and controlled in a functional organism [1, 2]. Despite many downstream rescue points where possible mistakes can be corrected [1, 3, 4], the initial mRNA readout of the maternal Bicoid gradient by the *hunchback* gene is remarkably accurate and reproducible between embryos [5, 6]: it is highly expressed in the anterior part of the embryo, quickly decreasing in the middle and not expressed in the boundary part. This precision is even more surprising given the very short duration of the cell cycles (6-15 minutes) during which the initial Bicoid readout takes place and the intrinsic molecular noise in transcription regulation [7–9].

Even though most of our understanding of transcription regulation in the fly embryo comes from studies of fixed samples, gene expression is a dynamic process. The process involves the assembly of the transcription machinery and depends on the concentrations of the maternal gradients [10]. Recent studies based on single-cell temporal measurements of a short lived luciferase reporter gene under the control of a number of promoters in mouse fibroblast cell cultures [11, 12] and experiments in *E. Coli* and yeast populations [13–16] have quantitatively confirmed that mRNAs are produced in bursts, which result from periods of activation and inactivation. What are the dynamical properties of transcription initiation that allow for the concentration of the Bicoid gradient and other maternal factors to be measured in these short intervals between mitosis?

In order to quantitatively describe the events involved in transcription initiation, we need to have a signature of this process in the form of time dependent traces of RNA production. Recently, live imaging techniques have been developed to simultaneously track the RNA production in all nuclei throughout the developmental period from nuclear cycle 11 to cycle 14 [17, 18]. In these experiments, an MS2 cassette is placed directly under the control of an additional copy of a proximal *hunchback* promoter. As the gene is transcribed, mRNA loops are expressed that bind fluorescent MCP proteins. Their accumulation at the transcribed locus gives an intense localized signal above the background level of unbound MCP proteins (Fig. 1C) [19]. By monitoring the living embryo, we obtain a time dependent fluorescence trace that is indicative of the dynamics of transcription regulation at the *hunchback* promoter (Fig. 1B, D and F).

However the fluorescent time traces inevitably provide an indirect observation of the transcription dynamics. The signal is noisy, convoluting both experimental and intrinsic noise with the properties of the probe: the jitter in the signal is not necessary indicative of actual gene switching but could simply result from a momentarily decrease in the recording of the intensity. To obtain a sufficient strong intensity of the signal to overcome background fluorescence, a long probe with a large number of loops is needed, which introduces a minimum buffering time (in the current experiments the minimal buffering time is 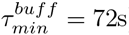) and preventing direct observation of activation [19].

**FIG. 1:**
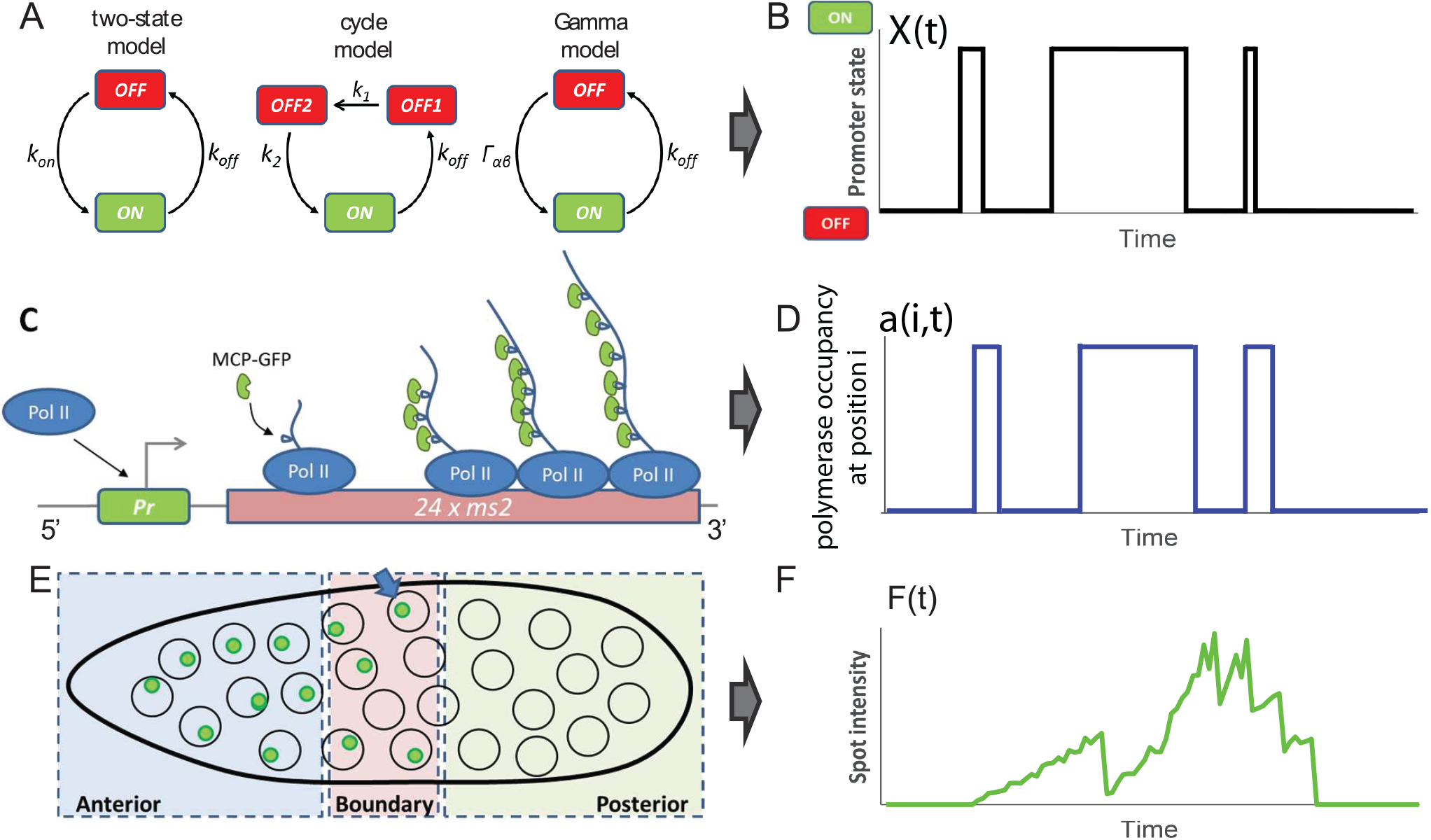
Transcription dynamics in the fly embryo. (A) The three models of transcription dynamics considered in this paper. From left to right: the two state model, the cycle model and the Gamma model (see SI Sections B, D and E). (B) Example of the promoter state dynamics (either ON or OFF) as a function of time. We assume that the polymerase is abundant and every time the promoter is ON and is not flanked by the previous polymerase a new polymerase will start transcribing. The black lines represent arrival times of the RNA polymerases to the promoter. (C) In the ON state, the promoter (*Pr*) is accessible to RNA polymerases (Pol II) that initiate the transcription of the target gene and the *24*× MS2 loops. As the *24*× MS2 mRNA is elongated MCP-GFP fluorescent molecules bind creating a detectable fluorescence signal. (D) The probability that site *i* on the gene was occupied by a polymerase as a function of time is given by the promoter occupancy in B and the finite size of the polymerase. (E) MCP-GFP molecules labeling several mRNA co-localize at the transcription loci, which appear as green spots under the confocal microscope. The spot intensities are then extracted over time and classified by each nuclei’s position in the *Drosophila* embryo as Anterior, Boundary and Posterior. (F) An example of the experimental signal: one spot's intensity a function of time, corresponding to the arrivals of RNA polymerases in (D) and the promoter state in (B).

To understand the details of the regulatory process that controls mRNA expression we need to quantify the statistics of the activation and inactivation times, as has been done in cell cultures [11, 12, 14, 15]. However the very short duration of the cell cycles (5-15 minutes for cell cycles 11-13) in early fly development prevents accumulation of statistics about the inactivation events and interpretation of these distributions. Direct observation of the traces suggests that, contrary to the previous reports[17, 18], transcription regulation is not static but displays bursts of activity and inactivity. However the eye can often be misleading when interpreting stochastic traces. In this paper we develop a statistical analysis of time dependent gene expression traces based on specially designed autocorrelation functions to investigate the dynamics of transcription regulation. This method overcomes the curse of naturally short traces caused by the limited duration of cell cycles that make it impossible to infer the properties of the regulation directly from sampling the activation and inactivation time statistics. Combining our analysis technique with models of transcription initiation and high resolution microscopy imaging of the MS2-MCP transgene under the control of the *hunchback* promoter, we show evidence suggesting that transcription initiation in cell cycles 12-13 is bursty. We focus on characterizing the transcription in the anterior and middle parts of the embryo and find that the dynamics is unchanged between cycle 12 and 13. We use these results to estimate the precision of the transcriptional readout. We show that the readout in each cell cycle is relatively imprecise compared to the precision of the mRNA measurement obtained on fixed samples [6].

## II. RESULTS

### A. Characterizing the time traces

We study the transcriptional dynamics of *hunchback* by generating embryos that express an MS2-MCP reporter cassette under the control of the proximal *hunchback* promoter (Fig. 1C), using previously developed techniques [17, 18], with an improved MS2 reporter [20] (see Materials and Methods for details). The MS2-MCP cassette was placed towards the 3’ end of the transcribed sequence and contained 24 MS2 loop motifs. While the gene is being transcribed, each newly synthesized MS2 loop binds a MCP-GFP molecule. In each nucleus, where transcription at this transgene is ongoing, we observe a unique bright fluorescent spot, which corresponds to the accumulation of several MS2-containing mRNAs at the locus (Fig. 1C). We assume that the fluorescent signal from a labelled mRNA disappears from the recording spot when the RNAP reaches the end of the transgene. With this setup we image the total signal in four fly embryos using confocal microscopy, simultaneously in all nuclei (Fig. 1E) from the beginning of cell cycle (cc) 11 to the end of cell cycle 13. We obtain a signal that corresponds to the temporal dependence of the fluorescence intensity of the transcriptional process in each nucleus, which we refer to as the time trace of each spot. Fig. 1F shows a cartoon representation of such a trace resulting from the polymerase activity (Fig. 1D) dictated by the promoter dynamics (Fig. 1B). We present examples of the traces analyzed in this paper in Fig. ?? and the signal preprocessing steps in the Materials and Methods and SI Section A.

To characterize the dynamics of the *hunchback* promoter we need to describe its switching rates between ON states, when the gene is transcribed by the polymerase at an enhanced rate and the OFF states when the gene is effectively silent with only a small basal transcriptional activity (Fig. 1A and B). Estimating the ON and OFF rates directly from the traces is problematic due to the high background fluorescence levels coming from the unbound MCF-GFP proteins that make it difficult to distinguish real OFF events from noise. To overcome this problem, we consider the autocorrelation functions of the signal. To avoid biases from differential signal strengths from each nucleus, we first subtract the mean of the fluorescence in each nucleus, *F*(*t*_*i*_) – 〈*F*(*t*_*i*_)〉 and then calculate the steady state connected autocorrelation function of the fluorescence signal (equivalent to a normalized auto-covariance), *C*(τ), at two time points separated by a delay time τ, *F*(*t*_*i*_) and *F*(*t*_*i*_ + τ), normalized by the variance of the signal over the traces, according to Eqs. 11 and 12 in Materials and Methods. We will always work with the *connected* autocorrelation function, which means the mean of the signal is subtracted from the trace. The autocorrelation function is a powerful approach since it averages out all temporally uncorrelated noise, such as camera shot noise or the instantaneous fluctuations of the fluorescent probe concentrations.

Fig. 2A compares the normalized connected autocorrelation functions calculated for the steady state expression in the anterior of the embryo (excluding the initial activation and final deactivation times after and before mitosis) in cell cycles 12 and 13 of varying durations: ~ 3 and ~ 6 minutes. The steady state signal from cell cycle 11 did not have enough time points to gather sufficient statistics. The functions decay as expected, showing a characteristic correlation time, then reaching a plateau at negative values before increasing again. Since the number of data points separated by large intervals is small the uncertainty increases with τ. Autocorrelation functions calculated for very long time traces have neither the negative plateau nor the increase at large τ. For example, the long-time connected autocorrelation functions shown in Fig. 2D calculated from the simulated trace of the process described in Fig. 1 and shown in Fig. 2C differ from the short time connected autocorrelation function in Fig. 2E calculated from the same trace (see SI Section G for a description of the simulations). As the traces get longer the connected autocorrelation function approaches the longtime results (Fig. 11) and the connected autocorrelation function of a finite duration trace of a simple correlated brownian motion (an Ornstein-Uhlenbeck process) displays the same properties (see Fig. 12). The dip is thus an artifact of the finite size of the trace. We also see that the autocorrelation functions shift to the left for short cell cycles (Fig. 2A), resulting in shorter correlation times, defined as the value of τ at which the autocorrelation function decays by *e*, for earlier cell cycles. However, calculating the autocorrelation functions for time traces of equal lengths for all cell cycles (Fig. 2B) shows that the shift was also a bias of the finite trace lengths, and after taking it into account, the transcription process in all the cell cycles has the same dynamics (although we note that the dynamics from this truncated trace is not the true long time dynamics).

This preliminary analysis shows that to extract information about the dynamics of transcription initiation we will need to account for the finite time traces. Additionally, a direct readout of even effective rates from the correlation time is difficult, because the autocorrelation due to the underlying gene regulatory signal (Fig. 1B) is obscured by the autocorrelation due to the timescale for the elongation of the sequence to be transcribed after the MS2 cassette (Fig. 1D) – the gene buffering time. The observed time traces are a convolution of these inputs (Fig. 1F). The form of the autocorrelation function and our ability to distinguish signal from noise also depends on the precise positioning and length of the fluorescent gene [19]. The analysis is thus limited by the buffering time of the signal, given as the length of the transcribed genomic sequence that carries the fluorescing MS2 loops divided by the polymerase velocity, and is only possible if the autocorrelation time of the promoter is larger than the buffering time. A construct with the MS2 transgene placed at the 3’ end of the gene (Fig. 4B) gives a reliable readout of the promoter activity even for fast switching between the two states but the weak signal is hard to distinguish from background fluorescence levels. Conversely, a 5’ positioning of the transgene (Fig. 4A) is insensitive to background fluorescence but can only be used to infer very slow switching [19].

**FIG. 2:**
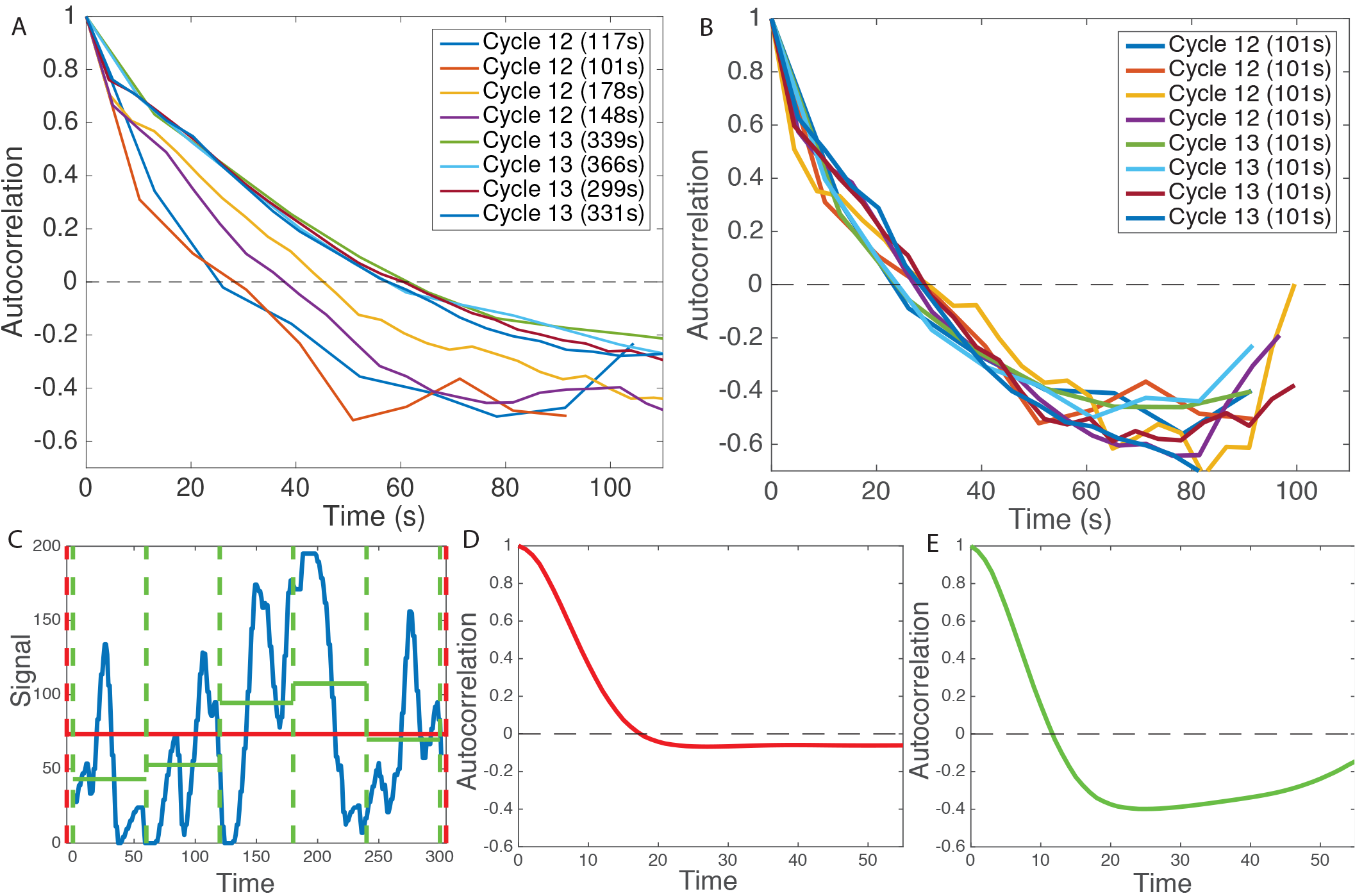
Autocorrelation analysis of fluorescent traces from cell cycles 12-13. (A) Autocorrelation functions for traces of different length caused by the variable duration of the cell cycle. Reading off the autocorrelation time as the time at which the autocorrelation function decays by a value of *e* would give different values for each trace. (B) Autocorrelation function calculated for the same traces reduced to have equal trace lengths, all equal to the trace length of the shortest trace, shows that the differences observed in panel A are due to finite size effects. (C) An example of a signal simulated for the process described in Fig.1 for a two state model for 300 seconds (blue). Taking the whole 300 second interval (red dashed) gives a good approximation of the average signal (red line) and the effect of finite size on the autocorrelation function is small (D). Reducing the time window to 60 seconds (green dashed line) correlates the average with the signal much more and the effect of finite size on the autocorrelation is strong (E). Parameters for the simulation in (C-E) are: *k*_on_ = *k*_off_ = 0.06*s*^-1^, sampling time *dt* = 4*s*, for the red curve *T* = 60*s* and *M* = 2000 nuclei, for the green curve *T* = 300*s* and *M* = 10000 nuclei (same total amount of data).

### B. Promoter switching models

The promoter activity we are interested in inferring can in principle be described by models of varying complexity (see Fig. 1A). In the simplest case, the gene is consecutively yet noisily expressed following a Poisson distribution of punctual ON events – this has previously been called a static promoter (not represented in Fig. 1A). Although the promoter dynamics would be uncorrelated in this case, the gene buffering would still produce a finite correlation time (see SI Section F). Alternatively, the promoter could have two well defined expression states: an ON state during which the polymerase is transcribing at an enhanced level and OFF state when it transcribes at a basal level. This situation can be modeled by stochastic switching between the two states with rates *k*_on_ and *k*_off_ (left panel in Fig. 1A and Materials and Methods). However, as was previously observed in both eukaryotic and prokaryotic cell cultures [11, 12, 14, 15], once the gene is switched off the system may have to progress through a series of OFF states before the gene can be reactivated. Recently these kinds of cycle models have been discussed for the *hunchback* promoter [21]. The intermediate states can correspond to, for example, the assembly of the transcription initiation complex, opening of the chromatin or transcription factor presence. These kinds of situations can either be modeled by a promoter cycle (middle panel in Fig. 1A and Materials and Methods), with a number of consecutive OFF states, or by an effective two state model that accounts for the resulting non-exponential, but gamma function distribution of waiting times in the off state (right panel in Fig. 1A and Materials and Methods). We present our method for all of these models and consider all but the gamma function distributed switching time model to learn about the dynamics of *hunchback* promoter dynamics.

### C. Autocorrelation approach

To infer the transcription dynamics from the data we built a mathematical model that calculates the autocorrelation functions that account for the experimental details of the probes, incorporating the MS2 loops at various positions along the gene and correcting for the finite length of the signal. The basic idea behind our approach is that while the initiation of transcription is stochastic and involves switching between the ON and possibly a number of OFF states (*X*(*t*) in Fig.1B denotes the binary gene expression state), the obscuring of the signal by the probe design is completely deterministic [18, 22], resulting in the probability *a*(*i*, *t*) that the polymerase is at position *i* at time *t* (Fig.1D). The promoter dynamics can thus be learned from the noisy autocorrelation function of the fluorescence intensity 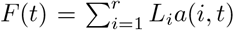 (Fig.1F), provided the parameters of the probe design encoded in the loop function *L*_*i*_ (positioning of the probe etc.) are known (Fig. 1C) and the signal is calibrated to know the fluorescence intensity coming from one loop [18].

Broadly, our model assumes that once the promoter is in an ON state the polymerase binds and deterministically travels along the gene producing MS2 loops containing mRNA that immediately bind MCP and result in a strong localized fluorescence (Fig. 3). We count the progression of the polymerase in discrete time steps, where one time step corresponds to the time of it takes the polymerase to cover a distance of 150 base pairs equal to its own length (Fig. 3A). The probability that there is a polymerase at position *i* at time *t*, *a*(*i*, *t*) is simply a delayed readout of the promoter state at time *t* − *i*, *a*(*i*, *t*) = *X*(*t* − *i*) where *t* is measured in polymerase time steps (Fig. 1B). We assume that polymerase is abundant and that at every time step a new polymerase starts transcribing, provided the gene is in the ON state (Fig. 1B and D). The amount of fluorescence produced by the gene at one time point is determined by the number of polymerases on the gene (Fig. 3A). The amount of fluorescence from one polymerase that is at position *i* on the gene depends on the cumulated number of loops that the polymerase has produced *L*_*i*_, where 1 ≤ *i* ≤ *r*, *r* corresponds to the maximum number of polymerases that can transcribe the gene at a given time and *L*_*i*_ = 1 corresponds to one loop fluorescing, as depicted in the cartoon in Fig. 3D. The known loop function *L*_*i*_ depends on the build and the position of the MS2 cassette on the gene, it is input to the model and does not necessarily take an integer value since the polymerase length and the loop length do not coincide (Fig. 3D). Given the steady state probability of the gene to be on *P*_on_ the average fluorescence in the steady state is:

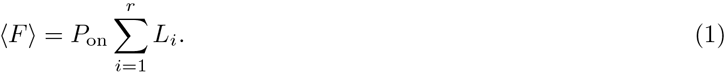

Since we assume the polymerase moves deterministically along the gene, seeing a fluorescence signal both at time *t* and position and *i* and at time *s* and position *j* means the gene was ON at time *t* − *i* and *s* − *j*, which is determined by how many loops (*i* and *j*) the polymerase has produced. Taking the earlier of these times, we need to calculate the probability that the gene is also ON at the later time. The autocorrelation function of the fluorescence can thus be written as:

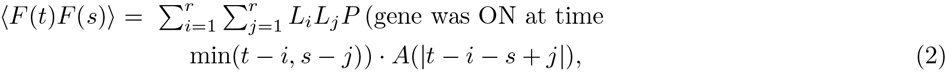

where *A*(*n*) is the probability that the gene is ON at time *n* given that it was ON at time 0. The precise form of *P*_on_, *P*(gene was on at timemin(*t* − *i*, *s* − *j*)) and *A*(|*t* − *i* − *s* + *j*|) depends on the type of the promoter switching model. We assume that the polymerase moves at constant speed along the gene and that there is no splicing throughout the transcription process. We give explicit expressions for all the models used in the Materials and Methods section and the Supplementary Information. Importantly, if we know the design of the construct, and calibrate the signal, we can use Eq. 1 to obtain the ratio of switching rates and Eq. 2 to obtain their particular values (see Materials and Methods).

**FIG. 3:**
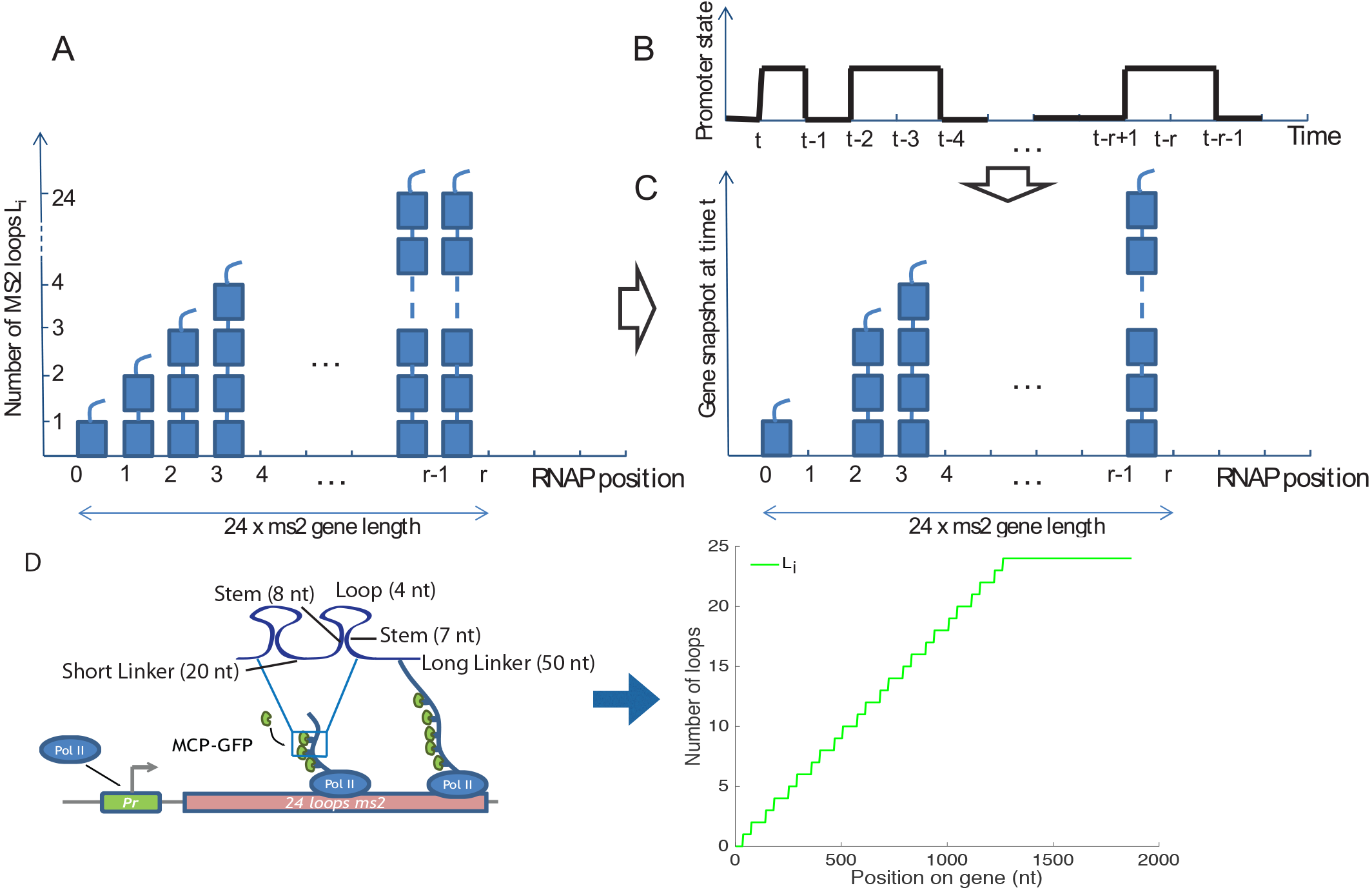
The gene expression model used in the autocorrelation function calculation. The autocorrelation inference approach is based on the idea that the stochastic transcriptional dynamics can be deconvoluted from the signal coming from the deterministic fluorescent construct, if we know the gene construct design. (A) A concatenation of snapshots of the gene from *r* consecutive time steps. A polymerase covers a length on the gene corresponding to its own length in one time step, producing one MS2 loop. The gene has total length *r* and at any position *i* along the gene *L*_*i*_ < 24 loops have been produced. (B) The promoter state as a function of time and (C) an instantaneous snapshot of the gene corresponding to transcription from this promoter. (D) The construct design is encoded in the loop function *L*_*i*_. As the polymerase moves along the gene it produces MS2 loops. *L*_*i*_ is an average representation in terms of polymerase time steps of how many loops have been produced by a single polymerase. It is based on the experimental design shown on the left of the panel.

To avoid biases coming from nucleus to nucleus variability, we calculated the normalized connected correlation function defined in Eqs. 11 and 12 in Materials and Methods. The theoretically calculated connected autocorrelation function, *C*_*r*_ (Eq. 13 which corresponds to the longtime correlation function in Fig. 2C and D) differs from the empirically calculated connected autocorrelation function from the traces, *c*(*r*) (Eqs. 11 and 12 in Materials and Methods, which corresponds to the short time correlation function in Fig. 2C and E) due to finite size effects coming from spurious correlations between the empirical mean and the data points. Since by definition the mean of a connected autocorrelation function is zero (see Eqs. 11 and 12 in Materials and Methods), the area under the autocorrelation function must be zero. For short traces this produces the artificial dip discussed in Fig. 2, which for long traces is not visible as it is equally distributed over long times. To compare our theoretical and empirical correlation functions we explicitly calculate the finite size correction and include this correction in our analysis (Materials and Methods and SI Section H and I).

In this paper, we have analyzed data from fly embryos with 3’ promoter constructs only, limiting ourselves to the steady state part of the trace. We limit our analysis to the steady state part of the interphase by taking a window in the middle of the trace to avoid the initial activation and final deactivation of the gene between the cell cycles (see Materials and Methods). However the method can also be applied to non-steady state systems (see SI Section C) and other constructs, including cross-correlation functions calculated from signals of different colors inserted at different positions along the gene (see SI Section J), which we discuss using simulated data.

### D. Simulated data

We first tested the autocorrelation based inference on simulated short-trace data with underlying molecular models with different levels of complexity for a construct with the MS2 probe in the 3’ end of the gene (Fig. 4B). In Fig. 4D we compare autocorrelation functions for the three state model for constructs with the MS2 loops positioned at the beginning of the transcribed region (5’, Fig. 4A) and at the end of the transcribed region (3’, Fig. 4B), and the cross-correlation function calculated from a two-colored probe construct (Fig. 4E). The analytical model correctly calculates the short trace autocorrelation function approach and is able to infer the dynamics of promoter switching for all models. It can also be adapted to infer the promoter switching parameters for any intermediate MS2 construct position, given of the limitations of each of the constructs discussed above [19].

The autocorrelation function based inference reproduces the underlying parameters of the dynamics with great accuracy for switching timescales smaller than the gene buffering time that obscures the signal (Fig. 4F). In Fig. 4F we show the results of the inference for the 3’ two state model for difference values of the ON and OFF rates, *k*_on_ and *k*_off_. For switching rates faster than the gene buffering time, the autocorrelation function coming from the length of the construct dominates the signal and the precision of the inference goes down. For very fast switching rates (> 0.12*s*^-1^), increasing the length of the traces or the number of nuclei (red vs blue curve above *k*_on_ + *k*_off_ = 0.1*s*^-1^ in Fig. 4F) does not help estimate the properties of transcription. For intermediate switching rates (0.07 – 0.12*s*^-1^), increasing the trace length or increasing the number of nuclei extends the inference range (black and green dashed lines vs blue solid line Fig. 4F) and in all cases increasing the number of nuclei decreases the uncertainty as can be seen from the smaller error bars (shown only for the red and blue lines for figure clarity).

Using two colored probes attached at different positions along the gene gives two measurements of transcription allows for an independent measurement of the speed of the polymerase – one of the parameters of the model that currently must be taken from other experiments. While the estimates of polymerase speed in the fly embryo are reliable [18], it has been pointed out as a confounding factor in other correlation analysis [23].

The autocorrelation approach also correctly infers the parameters of transcriptional processes when applied to traces that are out of steady state (see SI Section C). However, since the process is no longer translationally invariant more traces are needed to accumulate sufficient statistics. For this reason, in the current analysis of fly embryos we do not analyze the transient dynamics at the beginning and end of each cycle and we restrict ourselves to the middle of the interphase assuming steady state is reached.

### E. Fly trace data analysis

We divided the embryo into the anterior region, defined as the region between 0% and 35% of the egg length (the position at 50% of the egg length marks the embryo midpoint), where *hunchback* expression is high, and the boundary region, defined as the region between 45% and 55% egg length, where *hunchback* expression decreases. The mean probability for the gene to be ON during a given cell cycle *P*_on_ (restricted to the times excluding the initial activation and deactivation of the gene, which we will call the steady state regime), given by Eq. 1, is reproducible between the four embryos in cell cycle 12 and 13, both in the anterior region and at the boundary (Fig. 5A). The probability for the gene to be ON is over three fold higher in the anterior region than in the boundary and does not change with the cell cycle. *P*_on_ ~ 0.5 in the anterior indicates that in each nucleus the polymerase spends about half the steady state expression time transcribing the observed gene. At the boundary the gene is transcribed on average during about 10% of the steady state part of the cell cycle. The estimates for *P*_on_ in the earlier cell cycles were not reproducible between the four embryos, likely because the time traces were too short to gather sufficient statistics for this kind of analysis. We concentrated on cell cycle 12 and 13 for the remainder of the analysis.

**FIG. 4:**
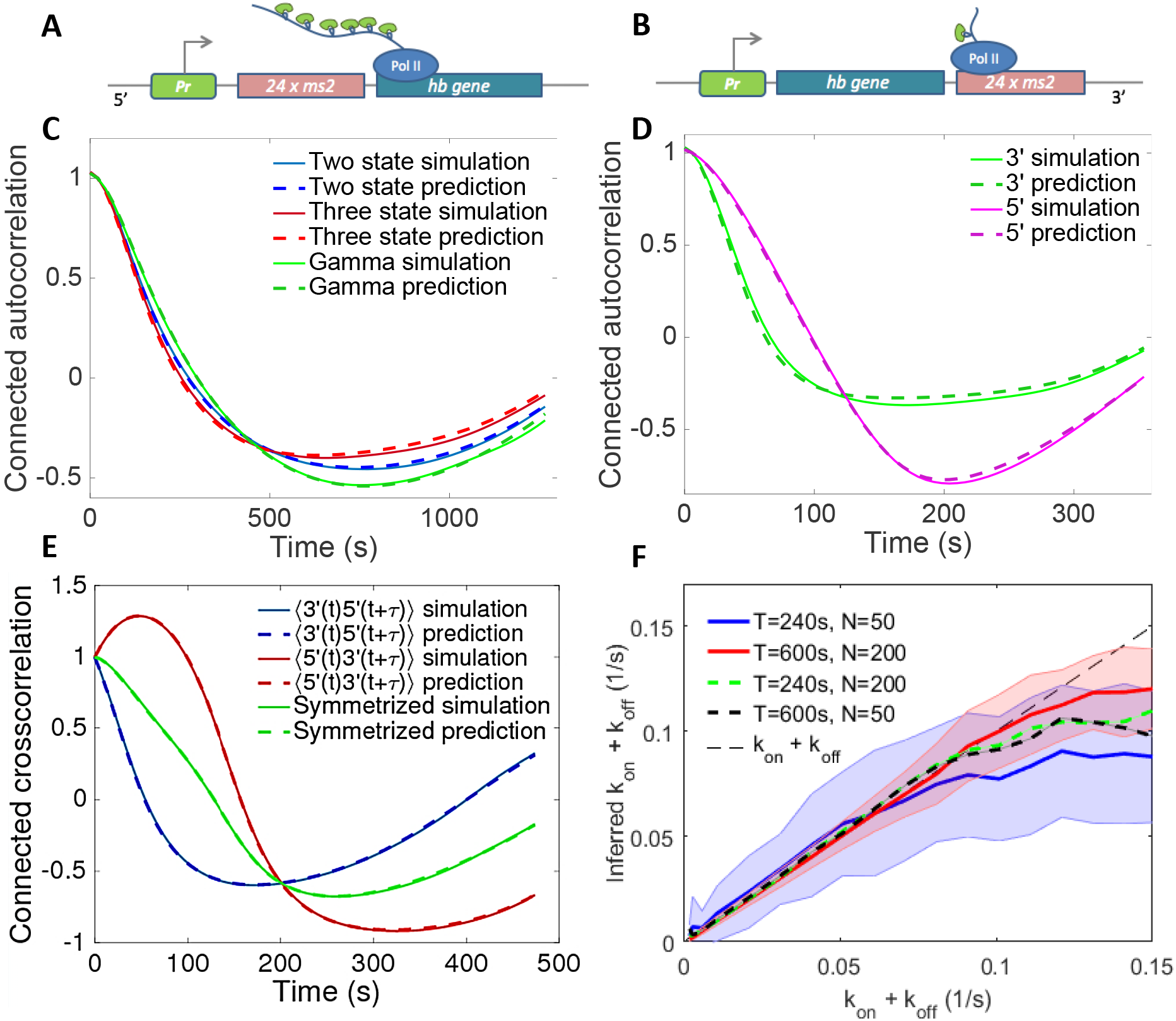
The autocorrelation based inference analysis performed on short trace simulated data for models of various complexity and positioning of the MS2 probe. Examples of the inferred autocorrelation functions fit to ones calculated from simulated traces (according to Gillespie simulations described in SI Section G) show perfect agreement for 3’ MS2 insertions assuming a two state (telegraph) model, three state model and gamma function bursty model (A), as well as 3’, 5’ for the two state model (B). C. The cross-correlation between the signal coming from two different colored fluorescent probes positioned at the 3’ and 5’ ends. D. The inference procedure for the two state model correctly finds the parameters of transcription initiation in a wide parameter range. The inference range grows with trace length and the number of nuclei. Error bars shown only for *T* = 240*s*, *N* = 50 nuclei (blue line) and *T* = 600*s*, *N* = 200 nuclei (red line) for clarity of presentation. Parameters for the simulations and predictions are: (C) For two state *k*_on_ = 0.005 s^-1^, *k*_off_ = 0.01 s^-1^, sampling time *dt* = 6 s, *T* = 360 s and number of cells *M* = 20000, for three state same parameters with *k*_off_ = 0.01 s^-1^, *k*_1_ = 0.01 s^-1^ and *k*_2_ = 0.02 s^-1^, for Γ model same parameters with *k*_off_ = 0.005 s^-1^ and *α* = 2 and *β* = 0.01 s^-1^. (D) *k*_on_ = 0.02 s^-1^, *k*_off_ = 0.01 s^-1^, sampling time *dt* = 6 s, *T* = 600 s and number of cells *M* = 20000. (E). *k*_on_ = 0.01 s^-1^, *k*_off_ = 0.01 s^-1^, *dt* = 6 s, *T* = 480 s and *M* = 20000. The 5’ construct is modeled as having 20 more fluorescent polymerase sites than the 3′ construct. F. *P*_on_ = 0.1

Based on the different behavior at the boundary and in the anterior, we separately inferred the transcriptional dynamics parameters in the two regimes, using the autocorrelation approach that corrects for finite time traces. The Poisson random firing model, the two and three state cycle models all provide reasonably good fits to the all the traces in both regions (see Fig. 5B for an example and Fig. 10 for the fits in both regions in all embryos). However, the fit of the Poisson random firing model (red line) only captures the short time behavior of the measured autocorrelation function. The two and three state model fits are indistinguishable and the two state fit is reproducible between cell cycles and embryos (Fig. 5B). The three state fit is reproducible at the level of the sum of the effective ON and OFF rates (same fit as shown for the two state model in Fig. 5C), but gives fluctuating values for *k*_1_/*k*_2_, the parameter determining how well it is approximated by a two state model (see Fig. 13, *k*_1_/*k*_2_ < 1 describes one fast reaction between the OFF states, effectively giving a two state model, while *k*_1_/*k*_2_ = 1 gives equal weights to the two reactions, clearly distinguishing two OFF states). Since the two state model is reproducible and has lesser complexity we will further consider the two state model.

**FIG. 5:**
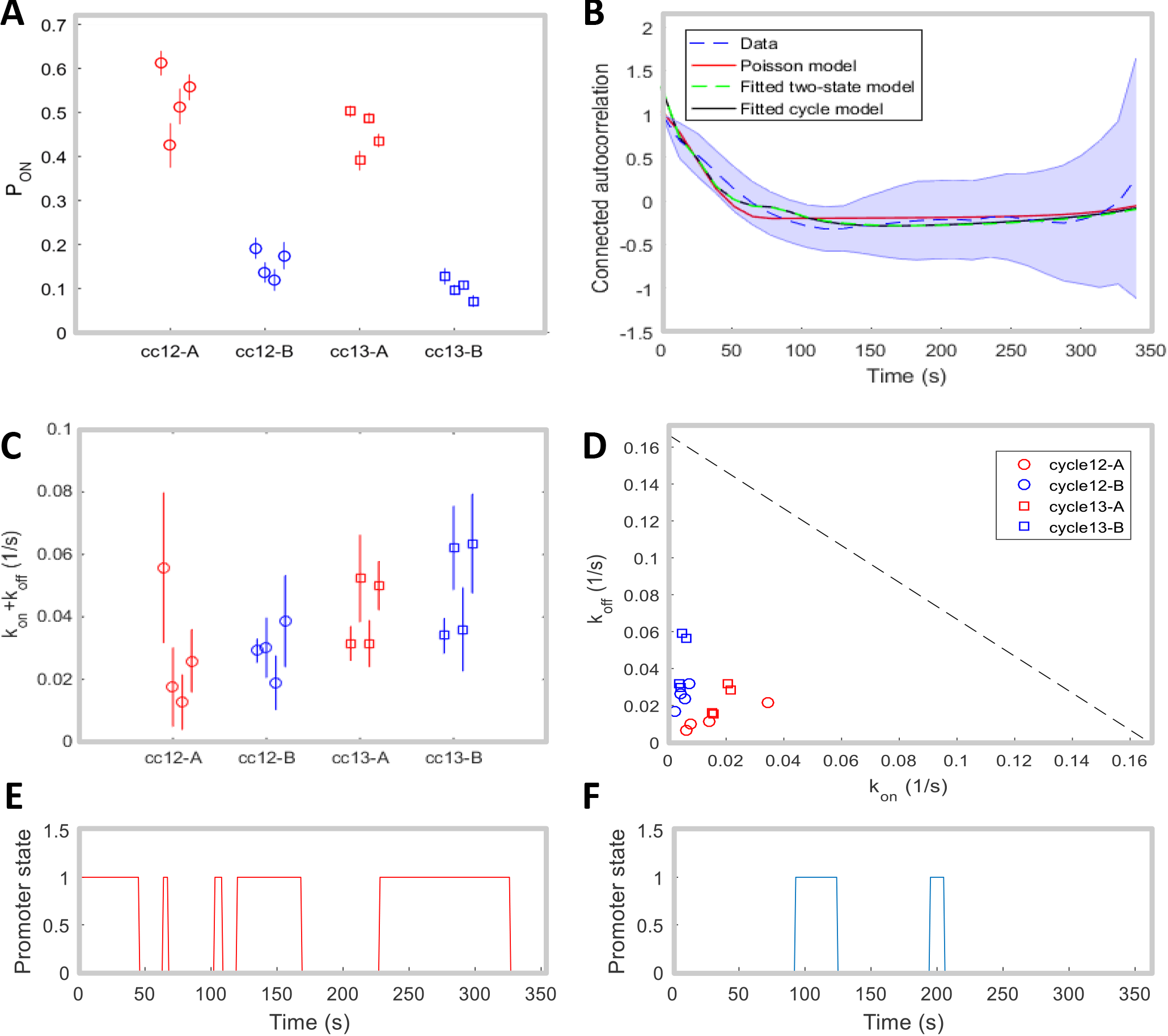
Inference results for fly data. (A) Inferred values of *P*_*ON*_ for different nuclei positions (A-Anterior, B-Bounary) and cell cycles. (B) Example of the mean connected autocorrelation function of the traces in cell cycle 13 (dashed blue line, with shaded error region) and of the fitted Poisson (red), two-state (green) and cycle (black) models. The fitted curves generated from the two-state and three state cycle model are almost superimposed. (C) Inferred values of *k*_on_ + *k*_off_ using the two-state model. In (A) and (C), the standard error bars are calculated by performing the inference on 20 random subsets that take 60% of the original data. (D) Inferred values of *k*_on_ and *k*_off_ in the Anterior (red) and Boundary (blue), in cell cycle 12 (circle) and cell cycle 13 (square). For each condition, 4 inferred values for 4 movies are shown. (E-F) Two trajectories of the promoter state with the inferred parameters in the Anterior (red) and Boundary (blue).

The inference procedure independently fits the characteristic timescale of the process, defined as the inverse of the sum of two rates, *k*_on_ + *k*_off_ (Fig. 5C), and then uses an independent fit of the probability of the gene to be ON, *P*_on_ (Fig. 5A), to disentangle the two rates (Fig. 5D). Examples of the promoter state over time with the rates' inferred values are shown in Fig. 5E (for the anterior region) and Fig. 5F (for the posterior region). Assuming the two state model we find that the characteristic timescale in most embryos is slighter shorter at the boundary (~ 25*s*) than in the anterior region (~ 33*s*) and the variability between the two cell cycles is comparable to the embryo to embryo variability (Fig. 5C). Both timescales are much larger than the 6s buffering time during which a second polymerase cannot bind because the first one has not cleared the binding site (shown as the gray dashed line in Fig. 5D), which sets a natural scale for the timescales we can infer. We find that in the anterior region of the embryo the two switching rates *k*_on_ and *k*_off_ show variability from embryo to embryo (between 0.009*s*^-1^ to 0.078*s*^-1^ – see Table I and II in the SI) but always scale together, which gives the observed one-half probability of the gene to be ON in a given nuclei during the steady state part of the interphase. Since the polymerase in the anterior on average spends half the steady state interphase window transcribing the gene, this suggests a clear bursting behavior of the transcription process, with switching between an identifiable active and inactive state of the promoter.

**FIG. 6:**
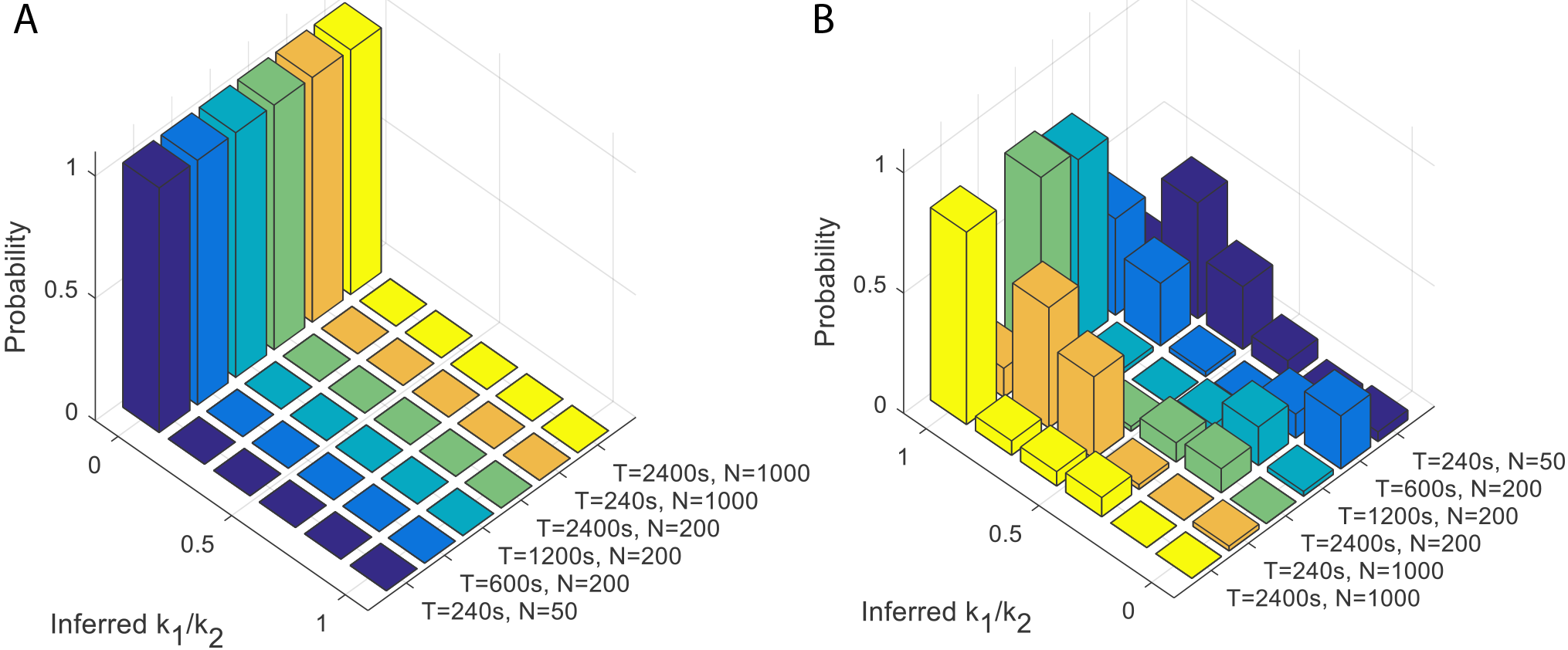
Longer time traces help distinguish between two state and three state cycle models. A. Inference on data generated by a two state model, which corresponds to *k*_1_/*k*_2_ = 0, from traces of different lengths *T* and using different numbers of nuclei *N* shows that longer traces help increase the probability to correctly learn the model type. Increasing the number of nuclei for short traces shows little improvement. The inference is repeated 50 times per condition. The experimental conditions studied in this paper are closest to the *T* = 240*s* and *N* = 50 nuclei panel. B. The same numerical experiment but assuming a three state cycle model, which corresponds to *k*_1_/*k*_2_ = 1. Parameters of the simulations: *P*_on_ = 0.1, *k*_off_ + 1/(1/*k*_1_ + 1/*k*_2_) = 0.02*s*^-1^ and *k*_1_/*k*_2_ = 0 in A, and *k*_1_/*k*_2_ = 1 in B.

*k*_on_ is much smaller at the boundary with very little embryo to embryo variability, while *k*_off_ has a similar range as in the anterior. This behavior is expected since high Bicoid concentrations in the anterior upregulate the transgene whereas lower concentrations at the boundary result in smaller activation rates. The ratio of the average *k*_on_ rates in the boundary and anterior is ~ 5, which can be compared to the 4 fold decrease expected from pure Bicoid activation, assuming the Bicoid gradient decays with a length scale of 100*μm* [24] and comparing the activation probabilities in the middle of the anterior and boundary regions. Given the crudeness of the this argument stemming from the variability of the Bicoid gradient in the boundary region and the uncertainty of the inferred rates, these ratios are in good agreement and suggest that a big part of the difference in the transcriptional process between the anterior and boundary is due to the change in Bicoid concentration. Of course other factors, such as maternal Hunchback, could also affect the promoter, leading to discrepancies between the two estimates.

The current data coming from four embryos and ~ 50 nuclei in each region with trace lengths of ~ 300*s* does not make it possible to distinguish between the two and three state models. We asked whether having longer traces or more nuclei could help us better characterize the bursty properties. We performed simulations with characteristic times similar to those inferred from the data (*k*_on_ + *k*_off_ = 0.01) assuming a two (Fig. 6A) and three state model (Fig. 6A). We then inferred the sum of the ON and OFF rates (*k*_on_ + *k*_off_) and the ratio of the two OFF rates (*k*_1_/*k*_2_). If the two OFF rates are similar (*k*_1_/*k*_2_ ~ 1) we infer a three state model. If one of the rates is much faster (*k*_1_/*k*_2_ ~ 0), we infer a two state model. We find that having more nuclei, which corresponds to collecting more embryos, would not significantly help our inference. However looking at longer traces would allow us to disambiguate the two scenarios, if the traces were 4 times longer, or ~ 20 minutes long. Since cell cycle 14 lasts for ~ 45 minutes, analyzing these traces could inform us about the effective structure of the OFF states. However in cell cycle 14, other genes get turned on after 15 minutes, so additional regulatory elements could be responsible for the observed transcriptional dynamics than in cell cycle 12 and 13. Our results suggest that with our current trace length we should be able to identify a two state model with large certainty, but we could not clearly identify a three state model. Our data may thus point towards a more complex model than two state, but a different kind of multistate model or a two state model obscured by other biases cannot be ruled out.

The error bars for the autocorrelation functions describe the variability between nuclei coming from both natural variability and measurement imprecision. While the autocorrelation function is insensitive to white noise, it does depend on correlated noise. The noise increases for large time differences τ, as the number of pairs of nuclei decreases and in our inference we reweigh the points according to their sampling so that the noise does not impair the precision of our inference. The error bars on the inferred parameter are due to variability between nuclei and are obtained from sampling different subsets of the data in each region and cell cycle. Additionally to the inter-nuclei and experimental noise there is natural variability between embryos. Since each nucleus transcribes independently and we assume similar Bicoid concentrations in each of the regions, the inter-embryo variability is of a similar scale as the inter-nuclei variability (Fig. 5C), as one expects given that the Bicoid gradient is incredibly reproducible between embryos [24].

### F. Accuracy of the transcriptional process

At the boundary, neighboring nuclei have dramatically different expression levels of the Hunchback protein. From measurements of the Bicoid gradient, Gregor and collaborators estimated that for two neighboring nuclei to make different readouts, they must be able to distinguish Bicoid concentrations that differ by 10% [25]. Following the Berg and Purcell [26] argument for receptor accuracy, and using measurements of diffusion constants for Bicoid proteins from cell cycle 14, the authors showed that, based on protein concentrations, the *hunchback* gene is not able to readout the differences in the concentrations of Bicoid proteins to the required 10% accuracy in the time that cell cycle 14 lasts. The authors invoked spatial averaging of Hunchback proteins as a possible mechanism that achieves this precision. Spatial averaging can increase precision, but it can also smear the boundary. Erdmann et al calculated the optimal diffusion constant Hunchback proteins must have for the averaging argument to work [27] and showed it is similar to experimental observations [5, 24]. However precision can already be established at the mRNA level and using static measurements Little and co-workers found that the relative variability of the mRNA transcribed from a *hunchback* locus in one nucleus is ~ 50% [6]. However measurements of cytoplasmic mRNA reduced this variability to ~ 10% [6].

Here we go one step further and use our direct measurements of transcription from the *hunchback* gene to directly estimate the precision with which the *hunchback* promoter makes a readout of its regulatory environment in a given cell cycle, *δP*_on_/*P*_on_. *δP*_on_/*P*_on_ is the relative error of the probability of the gene to be ON averaged over the steady state part of a cell cycle. Since the total number of mRNA molecules produced in a given cycle is proportional to *P*_on_ (shown in Fig. 14E as a function of embryo length), the precision at the level of *produced* mRNA in a given cycle is equal to the precision in the expression of the gene, *δ*mRNA/mRNA = *δP*_on_/*P*_on_. The accuracy of transcription activation is encoded in the stochasticity of gene activation. The gene randomly switches between two states: active and inactive, making a measurement about the regulatory factors in its environment and indirectly inferring the position of its nucleus. Since no additional information is provided by a measurement that is strongly correlated to the previous one, the cell can only base its positional readout on a series of independent measurements. Two measurements are statistically independent if they are separated by at least the expectation value of the time τ_*i*_ it takes the system to reset itself:

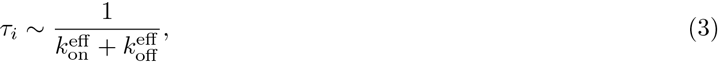

where in a two state model 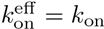 and 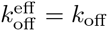. A more detailed estimate obtained by computing the variance of the time spent ON by the gene during the interphase (see SI Section K) shows that Eq. 3 underestimates the time needed to perform independent measurements. We find that for a two state model the accuracy of the readout of the total mRNA produced is limited by the variability of a two state variable divided by the estimated number of independent measurements within one cell cycle:

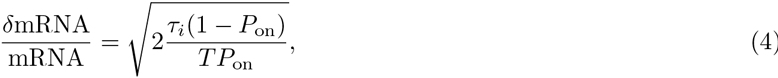

where *T* is the duration of the cell cycle and the factor 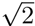 is a prefactor correction to the naive estimate. Eq. 4 is valid in the limit of *T* >> τ_*i*_ (the exact result if given in SI Section K). Using the rates inferred from the autocorrelation analysis (Fig. 5D) we see that the precision of the gene readout is much lower at the boundary than in the anterior, does not change with the cell cycle and is reproducible between embryos (ordinate in Fig. 7A). In the anterior part of the embryo it reaches ~ 50%, while at the boundary, it is very large, ~ 150%, even at cell cycle 13.

We can compare these theoretical estimates with direct estimates of the relative error of the total mRNA produced during a cell cycle, *δ*mRNA/mRNA, from the data. We divide the embryo into anterior and boundary strips, as we did for the inference procedure and calculate the mean and variance of *P*_on_. These empirical estimates of the precision of the gene measurement calculated agree with the theoretical estimates (Fig. 7A). We verified that our conclusions about the scale of our empirical estimates do not depend on the definition of the boundary and anterior regions (Fig. 14B). To see whether integrating the mRNA produced can increase precision we compared the empirical estimate of the steady state mRNA production (red line in Fig. 7B) to the relative error of the total mRNA produced in cell cycle 13 (blue line in Fig. 7B) and the total mRNA produced from cell cycle 10 to 13 (green line in Fig. 7B) averaged over embryos. We assumed that each nuclei has the total mRNA produced in cell cycle 13, 1/2 of the total mRNA produced by its mother in cell cycle 12, 1/4 of the mRNA produced by its grand-mother in cell cycle 12 etc. While we see about a 1/3 increase in the precision at the boundary from integrating the mRNA produced in different cell cycles, the estimate in the anterior region is not helped by integration over the cell cycles.

Since we are not able to rule out the three state cycle model as an accurate description of the transcriptional dynamics, we calculated the relative error assuming the same 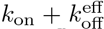 for a three state cycle 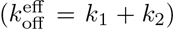 as for a two state model 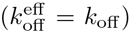 for different values of *k*_on_ and 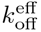 (Fig. 7C). We found that the relative error is always lower for the three state cycle model and the error decreases, regardless of the duration of the cell cycle, and as expected from Eq. 4 as the relative error is decreased by increasing *k*_on_ and decreasing *k*_off_. However the increase in precision from a three state cycle model in the parameter regime we inferred from the fly embryo is relatively modest.

Many previous analysis of precision from static images calculated the relative error of the distribution of a binary variable, which in each nucleus was 1 if the nucleus expressed mRNA in the snapshot, and 0 if it did not express[28, 29]. We analyzed our data using this definition of activity (see Fig. 14D for mean activity as a function of position) and found that for most embryos the relative error in the anterior drops to zero (Fig. 14C), indicating that all nuclei in a given region show the same expression state, but at the boundary the precision is still ~ 50%, in agreement with previous reports about the total mRNA in the nucleus [6]. This provides additional evidence for the bursty nature of transcription in the anterior of the embryo.

## III. DISCUSSION

Contrary to initial reports [17, 18] about the static nature of transcription initiation controlled by the *hunchback* promoter in fly development we show that the promoter is bursty with distinct periods of enhanced polymerase transcription followed by identifiable periods of basal polymerase activity. Our conclusions are based on a new autocorrelation based analysis approached applied to live imaging MS2-MCP data. The data we used in this paper was generated with a modified MS2 cassette [20] compared to the previously published data [17]. However the difference in our conclusions mainly comes from a detailed analysis of the traces.

Quantification of transcription from time dependent fluorescent traces in prokaryotes and mammalian cell cultures has shown that the promoter states cycle through at least three states [11, 12]. In one of these states the polymerase transcribes at enhanced levels, while in most of the remaining states the transcription machinery gets reassembled or the chromatin remodels. We find that in the anterior part of a living developing fly embryo, the *hunchback* promoter also cycles through at least two states, although we cannot conclusively rule out the possibility of more states when the gene is inactive. The main impediment to distinguishing different types of transcriptional cycles comes from the very short durations of the interphase in the early cell cycles when the *hunchback* gene is expressed. We showed that increasing the number embryonic samples would not help us distinguish between two and three state models, however looking at longer time traces would be informative (Fig. 6). Since cell cycle 14 lasts about 45 minutes, our analysis shows that the steady state part of the interphase provides enough time to gather statistics that can inform us about the detailed nature of the bursts. Unfortunately, other transcription factors such as the other gap genes regulate *hunchback* expression in cell cycle 14, possibly changing the nature of the transcriptional dynamics in a time dependent manner. We showed that the transcriptional dynamics is constant and reproducible in the earlier cell cycles (12-13) (Fig. 2), so independently of the question of the nature of the bursts it would be very interesting to see whether and how it changes when the nature of regulation changes.

Alternatively to looking at longer traces, a construct with two sets of MS2 loops placed at the two ends of the gene that bind different colored probes could be used to learn more about transcription dynamics [30]. We do not have access to data coming from such a promoter, but our analysis approach can be extended to calculate the cross-correlation function between the intensities of the two colored probes. Such cross-correlation analysis have previously been used to study transcription in cell cultures [31], transcriptional noise [32] and regulation in bacteria [33, 34]. Our theoretical prediction for such a cross-correlation function agrees with simulation results (Fig. 4C). Unfortunately, the cross-correlation function with one set of probes inserted at the 5’ end and the other at the 3’ shares the same problems of a 5’ construct. For fast switching rates, such a cross-correlation function suffers from the large buffering time (~ 300*s* in[18]) drawback of the 5’ design and can only be used for inferring large switching rates [35]. However, it does gives us access into dynamical parameters of transcription such as the speed of polymerase and it is able to characterize whether mRNA transcription is in fact deterministic and identify potential introns. Possibly, cross-correlations from two colored probes both inserted closer to the 3’ end could be optimal designs.

**FIG. 7:**
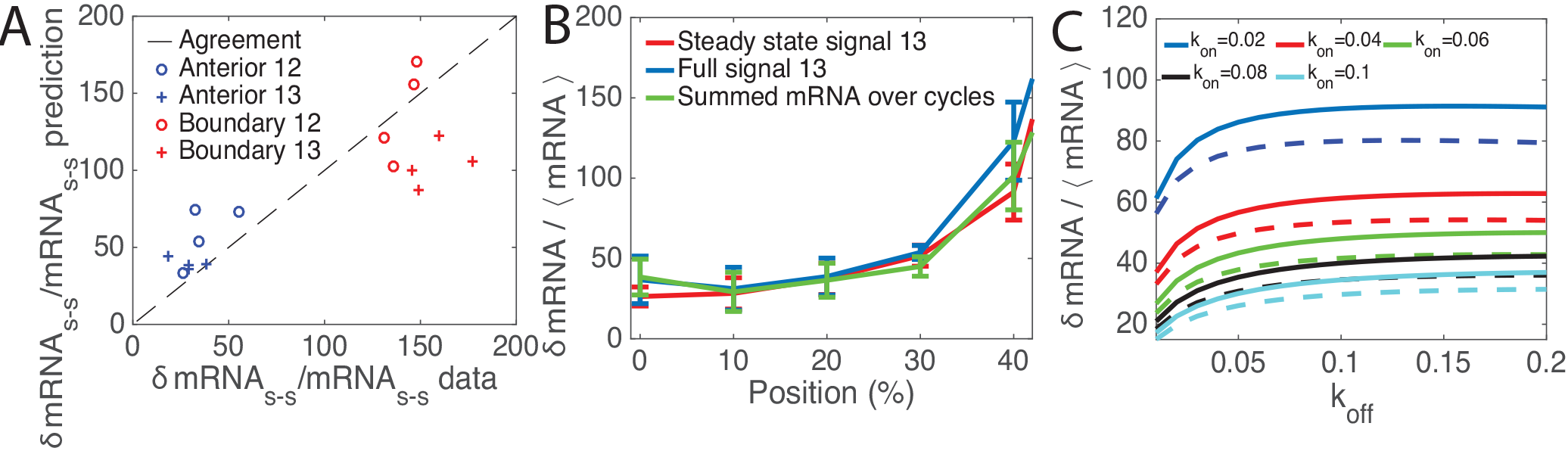
Precision of the *hunchback* gene transcription readout. A. Comparison of the relative error in the mRNA produced during the steady state of the interphase estimated empirically from data (abscissa) and from theoretical arguments in Eq. 4 using the inferred parameters in Fig. 5C (ordinate), in the anterior (blue) and the boundary (red) regions, show very good agreement. B. The relative error in the total mRNA produced in cell cycle 13 directly estimated from the data as the variance over the mean of the steady state mRNA production (red line, same data as in A), sum of the intensity over the whole duration of the interphase (blue line) and the total mRNA produced during cell cycles 11 to 13 (green line) for equal width bins equal to 10% embryo length at different positions along the AP axis. Each line describes a an average over four embryos (see Fig. 14C for the same data plotted separately for each embryo) and the error bars describe the variance. To calculated the total mRNA produced over the cell cycles, we take all the nuclei within a strip at cell cycle 13 and trace back their lineage through cycle 12 to cycle 11. We then sum the total intensity of each nuclei in cell cycle 13 and half the total intensity of its mother and 1/4 of its grandmother. C. Comparison of the relative error in the mRNA produced during the steady state for a two state, *k*_1_/*k*_2_ = 0, (solid lines) and three state cycle model, *k*_1_/*k*_2_ = 1, (dashed lines) with the same 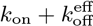 for different values of *k*_on_ and *k*_off_ shows that the three state cycles system allows for greater readout precision.

We assumed an effective model that describes the transcription state of the whole gene and does not explicitly take into account the individual binding sites. As a result all the parameters we learn are effective and describe the overall change in the expression state of the gene and not the binding and unbinding of Bicoid to the individual binding sites. For concreteness we presented our model assuming a change in the promoter state and constitutive polymerase binding, but our current model does not discriminated between situations where the transcriptional kinetics are driven by polymerase binding and unbinding and promoter kinetics. The presented formalism can be extended to more complex scenarios that describe the kinetics of the individual binding sites and random polymerase arrival times. Since we already have little resolution power to discriminate between these effective models, we chose to interpret the results of only these effective models. The exact contribution of the individual transcription binding sites could be inferred from the activity of promoters with mutated binding sites.

The time traces we had to analyze are very short and finite size effects are pronounced. Unlike in cell culture studies, where long time traces are available, we could not collect enough ON and OFF time statistics to characterize the promoter dynamics from the waiting time distributions. In this paper we show that simple statistics, the auto-and cross-correlation functions are powerful general tools that can be used in these kinds of challenging circumstances.

The approach we propose is a general method that can be used for any type of time trace analysis. However it becomes very useful in studying in vivo biological process where the biology naturally limits the available statistics. In our case the number of ON and OFF events is naturally limited by the short duration of the cell cycles. Our method explicitly calculates correlation functions for short traces, correcting for the finite size effects, and can be also used without making steady state assumptions about the dynamics (although this requires collecting sufficient statistics about two time points, which may be hard for short traces). With these corrections we see that while an effective two state model of the underlying dynamics of transcription regulation holds in the anterior and boundary regions of the embryo in all of the early cell cycles, the rates are different in the boundary and anterior regions, showing a strong dependence on position dependent factors such as Bicoid or maternal Hunchback concentrations. More statistics will make it possible to build more explicit models of Bicoid dependent activation.

In all cases, the rates that we can infer from time dependent traces are naturally limited by the timescales at which the polymerase leaves the promoter, which in our case is estimated to be ~ 6*s*. If the switching rates are faster than this scale, even a perfect, noiseless and infinitely accurate sampling of the dynamics will not be able to overcome this natural limit.

Our method requires knowing the design of the experimental system (number and position of the loops), the speed of polymerase as input and calibrating the maximal fluorescence from one gene. Measurements using two colored probes positioned at a distance on the same gene combined with a cross-correlation function analyses could access parameters such as the speed of polymerase and verify assumptions about the monotonous progression of the polymerase. Such effects can easily be incorporated into the model. While the polymerase speed is an important parameter and erroneous assumption could influence the inference, we have shown that our inference is relatively insensitive to polymerase speeds (see Fig. 15). In the current experiments we do not have an independent calibration of the maximal fluorescence coming from one gene, which could introduce potential errors in our analysis. However the reproducibility of our results suggests that these potential errors are small.

The presented analysis is an investigation of transcription dynamics from time dependent traces in living functioning organisms. It shows that the functional promoter that controls the first regulatory steps in fly development is bursty, even in the region with the highest activator concentration. The inferred rates are reproducible between nuclei and embryos and the inter embyo variability is similar to the inner embryo variability (Fig. 5A, C and D).

We used the obtained results to estimate the precision of the transcriptional process from the *hunchback* promoter. We found that even in the boundary region the variability in the mRNA produced in steady state by the different nuclei is large, with a relative error of about 50% (Fig. 7A). This variability further increases to 150% of the mean mRNA produced at the boundary. These empirical estimates are completely explained by theoretical arguments that treat the gene as an independent measuring device that samples the environment, correcting for the number of independent measurements during a cell cycle. In both cases, the precision at the level of the gene readout is not sufficient to form the precise Hunchback boundary up to half a nuclear width [36]. However, although we can extend our argument to the total mRNA produced in the early cell cycles (Fig. 7B), we do not know the amount of maternal *hunchback* mRNA in the nuclei. Having an irreversible promoter cycle could increase the theoretical precision, but only slightly in the parameter regime we have inferred and it would not change the quantitative conclusions about low precision backed by the empirical results.

In the same spirit, the construct we used here was limited to the 500 bp of the proximal *hunchback* promoter, which recapitulates the formation of a sharp boundary at later cell cycles in Fluorescent In Situ Hybridization (FISH) [20]. It is possible that the boundary phenotype is recovered thanks to averaging of mRNAs and proteins produced by the real gene or the transgenes in other nuclei. In the latter case, this would point towards a robust ‘safety” averaging mechanism that relies on the population. Alternatively, we have to be aware that the sharp boundaries were only detected on fixed samples and that having access to the dynamics of the transcription process likely provides a more accurate view on the process. We calculated and estimated from the data the precision of the gene readout based on the variability of the transcription process between nuclei. We find that the transcriptional process at a given position is quite noisy. Previous estimates of precision were based on static data and did not consider the probability of the gene to be ON, but assumed a binary representation where each nuclei is either active or inactive. By analyzing the full dynamic process we show that the gene is bursty and the transcriptional process itself is much more variable. Reducing the information contained in our traces to binary states, we find precise expression in the anterior, but still large variability at the boundary, similarly to previous results from Fluorescent In Situ Hybridization (FISH)[6].

Assuming that the precision in determining the position of the nuclei is encoded in the precision of the gene readout, a gene with the dynamics characterized in this paper needs to measure the signal ~ 200 times longer at the boundary to achieve the observed ~ 10% precision. A gene in the anterior would need to integrate only ~ 25 times longer. These results again suggest that the precision in determining the position of the nuclei is not only encoded in the time averaged gene readout, but probably relies either on spatial averaging mechanisms [25, 27, 37] or more detailed temporal information.

In summary, the early developing fly embryo provides a natural system where we can investigate in a functional setting the dynamics of transcription in a living organism. In our data analysis we are confronted by the same limitations that natural genes face: an estimate of the environmental conditions must be made in a very short time. Analysis of dynamical traces suggests that transcription is a bursty process with relatively large inter-nuclei variability, suggesting that simply the templated one to one time-averaged readout of the Bicoid gradient is unlikely. Comparison of mutant experiments can shed light on exactly how is the decision to form the sharp *hunchback* mRNA and protein boundary made.

## IV. MATERIALS AND METHODS

## V. EXPERIMENT PROCEDURE

### A. Constructs

For live monitoring of *hb* transcription activity in Drosophila embryos, we used the MS2-MCP system which allows fluorescent labeling of RNAs as they are being transcribed [17, 35, 38]. To implement the reporter system in embryos, we generated flies transgenic for single insertions of a P-element carrying *hb* proximal promoter upstream of an iRFP-MS2 cassette carrying 24 MS2 repeats [17, 39]. The flies also carry the P{mRFP-Nup107.K} [40] transgenic insertion on the 2^*nd*^ chromosome and the Pw[+mC]=Hsp83-MCP-GFP transgenic insertion on the 3rd chromosome. These allow the expression of the Nucleoporin-mRFP (mRFP-Nup) for the labeling of the nuclear envelopes and the MCP-GFP required for labeling of nascent RNAs [38]. All stocks were maintained at 25°C.

### B. Live Imaging

Embryo collection, dechorionation and imaging have been done as described in [17]. Image stacks (~19Z × 0.5*μ*m, 2*μ*m pinhole) were collected continuously at 0.197*μ*m XY resolution, 8bits per pixels, 1200x1200 pixels per frame. A total of 4 movies capturing 4 embryos from nuclear cycle 10 to nuclear cycle 13 were taken. Each movie, due to having different scanned field along the embryos' width, has a different time resolution: 13.1 *s*, 10.2 *s*, 5.1 *s* and 4.3 *s*.

### C. Image analysis

Nuclei segmentation, tracking and MS2-MCP loci analysis were performed as in [17] and recapitulated here. All steps were inspected visually and manually corrected when necessary. Nuclei segmentation and tracking were done by analyzing, frame by frame, the maximal Z-projection of the movies' mRFP-Nup channel. Each image was fitted with a set of nuclei templates, disks of adjustable radius and brightness comparable with raw nuclei's, from which the nuclei positions are extracted. During the cycle's interphase, each nucleus was tracked over time with a simple minimal distance criterion. For the analysis of MS2-MCP loci detection and fluorescent intensity quantification, the 3D GFP channel (MS2-MCP) were masked with the segmented nuclei images obtained in the previous step. This procedure also helps associating spots to nuclei. We then applied a threshold equal to 2 times the background signal to the masked images and selected only the connected regions with an area larger than 10 pixels. The spot positions are set as the position of the centroids of the connected regions. The intensity of each spot was calculated by summing up all the pixel intensity in the vicinity of the centroids (region of 1.5*μ*m × 1.5*μ*m × 1*μ*m) subtracted to the background intensity extracted from the region around and excluding the spots. In the (rare) case of multiple spots detected per nucleus, the biggest spot was selected.

For each nucleus, we collected the nucleus’ position and the spot intensity over time (here referred as ‘traces”). The traces were then classified according to their respective embryos (out of 4 embryos), cell cycle (10 to 13) and position along AP axis (either Anterior or Boundary). See SI Section A and Fig. 8 for examples of traces.

### D. Trace preprocessing

Before the autocorrelation function can be calculated the traces need to be preprocessed. To ensure that the data captures the dynamics of gene expression in its steady state, for each embryos and each cell cycle, we observed the spot intensity only in a specific time window. The beginning and the end of this window is determined as the moment the mean spot intensity over time of all traces (both at the anterior and the boundary) reaches and leaves its an expression plateau (see example in Fig. 9).

### E. The two state model

The detailed form of the autocorrelation function in Eq. 2 depends on the underlying gene promoter switching model. For the two state – telegraph switching model (left panel in Fig. 1A) the jumping times between the two states are both exponential and the dynamics is Markovian. The mean steady state probability for the promoter to be ON is *P*_on_ = *k*_on_/(*k*_on_ + *k*_off_), which combined with Eq. 1 gives the form of the mean fluorescence 〈*F*〉. The probability that the gene is ON at time *n* given that it was on at time 0 is *A_n_* = *P*_on_ + *e*^(*δ*-1)*n*^(1 − *P*_on_), where *δ* =1 − *k*_on_ − *k*_off_. The steady state connected correlation function depends only on the time difference (see SI Section B):

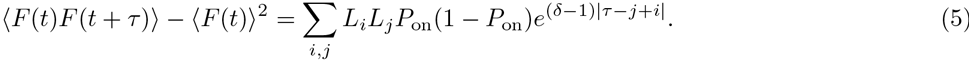

### F. The cycle model

In the cycle model (center panel in Fig. 1A) the OFF period is divided into different sub-steps that correspond to *K* intermediate states with exponentially distributed jumping times from one to the next. The transition matrix *T* encodes the rates of this irreversible chain. The probability of the promoter to be in the ON state is:

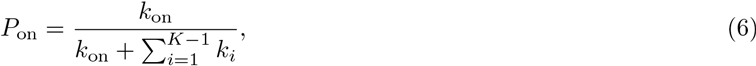

and that the steady state connected autocorrelation at is (see SI Section D):

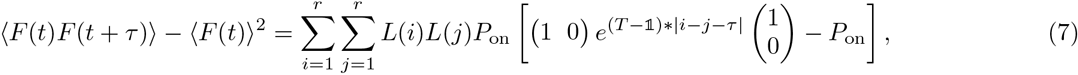

where τ is counted in polymerase steps. In the simple case of a two state model Eq. 7 reduces to Eq. 5.

### G. The γ waiting time model

An alternative description of a promoter cycle relies on a reduced description to an effective two state model where we use the fact that the transitions between the states are irreversible. The distribution of times spent in the effective OFF state τ, is no longer exponential, as it was in the two state model, but it has a peak at nonzero waiting times, which can be approximated by a Gamma distribution

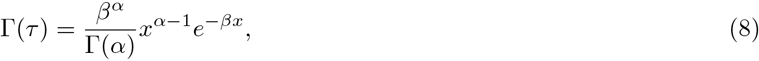

with mean *α/β* where *β* is the scale parameter, *α* is the shape parameter and Γ(*α*) is the gamma function. The true distribution of waiting times in a cycle model approaches the γ distribution if the OFF rates are all the same and *k*_off_ << 1. In this limit *β* ≈ *k*_off_, and *α* describes the number of intermediate OFF states. In the more general case it correctly captures the effective properties of the process. The mean probability of the promoter to be in ON state in the γ waiting time model is given by

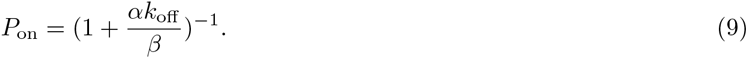

The autocorrelation function cannot be computed directly analytically. The steady state Fourier transform of the steady state autocorrelation is (see SI Section E):

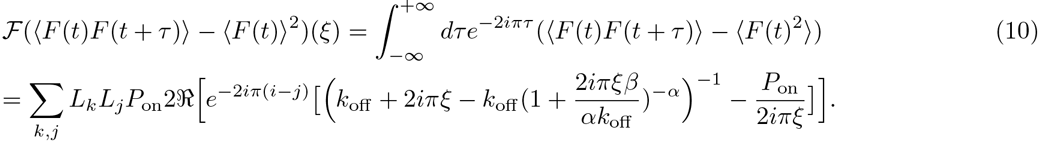

### H. Finite cell cycle length correction to the connected autocorrelation function

Due to the short duration of the cell cycle, the theoretical connected correlation functions need to be corrected for finite size effects when comparing them to the empirically calculated correlation functions. When analyzing the data we calculate the autocorrelation function from *M* traces {*v_α_*}_1≤*α*≤*M*_ of the same length *K, v_α_* = {*v_αj_*}_1≤*j*≤*K*_. We calculate the connected autocorrelation function for each trace and normalize it to 1 at time *t* = 0 to avoid spurious nucleus to nucleus variability:

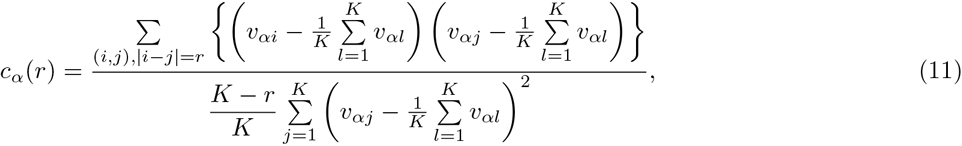

and then average over all *M* traces to obtain the final connected autocorrelation function:

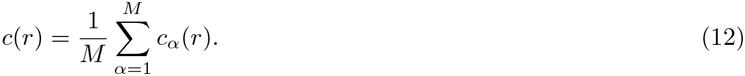

For 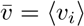 – the steady state true theoretical average of the random fluorescence intensity over random realization of the process, and 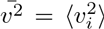 – the true theoretical second moment of the fluorescence signal, when *K* → ∞ the average over time points is equal to the theoretical average, 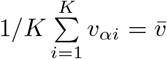 and the using time invariance in steady state the autocorrelation function becomes:

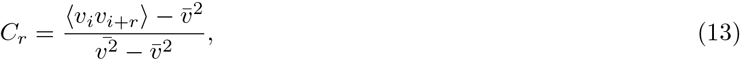

where 〈·〉 is an average over random realizations of the process. Eq. 13 corresponds to the limit we calculated in the theoretical model. To account for the finite size effects that arise due to short time traces we need to correct for the fact that for short traces 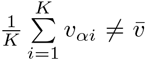 and 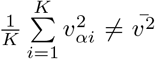 but both the mean and the variance are functions of *K*.

We note that for short traces the definitions of autocorrelation and autocovariance differ:

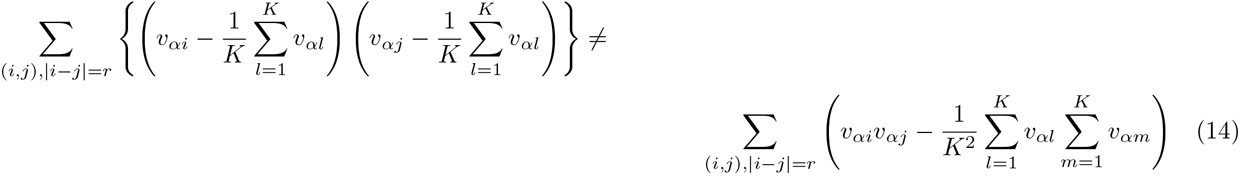

In practice for the analyzed dataset we found that the finite size effects for the variance can be neglected, however the mean over time points is a bad approximation to the ensemble mean. We present the finite size correction to the mean below. For completeness we include the finite size correction for the variance in SI Section I, although we do not use it in the analysis due to its numerical complexity and small effect.

If the variance of the normalized fluorescence intensity over random realizations of the process is well approximated by the average over the *K* time points, we can replace the denominator in Eq. 11 by 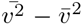 and in steady state evaluate the mean connected autocorrelation function (see SI Section H for details):

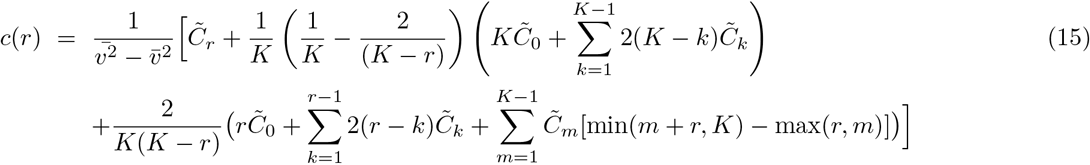

where 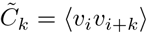 is the theoretical steady state non-connected correlation function of the process and the average is over random realizations of the process. If *v_i_* = *X*(*i*) then *C_k_* is proportional to *A*(*k*).

### I. Inference

The inference proceeds in three steps:

Step 1. Signal calibration. The intensity of the measured signal depends on a constant trace dependent offset value 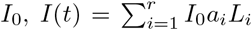. To calibrate this offset we take the maximum expression to be the mean of the maximun expression over all traces in a given region 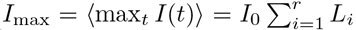. The calibrated fluorescence signal used in the analysis is then 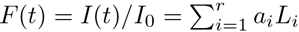. *P*_on_ is directly calculated using Eq. 1.

Step 2. Estimating parameter ratios. The ratios of the rates can be estimated directly from the steady state mean fluorescence values using Eqs. 6 and 9.

Step 3. Estimating parameters. Using the estimate for the ratio of the rates, the ON and OFF rates are found by minimizing the mean squared error between the data and the model.

## Acknowledgments

This work was supported by a Marie Curie MCCIG grant No. 303561(AMW), PSL IDEX REFLEX (ND, AMW, MC), ARC PJA20151203341 (ND), ANR-11-LABX-0044 DEEP Labex (ND), ANR-11-BSV2-0024 Axomorph (ND and AMW) and PSL ANR-10-IDEX-0001-02.

## VI. SUPPLEMENTARY INFORMATION: AUTOCORRELATION ANALYSIS

### A. Basic setup and data preprocessing

The raw data produced experimentally is a fluorescent signal *I*(*t*) measured at discrete times corresponding to the sampling time frame of the movie (see SIFig. 8 for examples of traces). At each locus and at each time point it is the sum of the background signal and a number of fluorescent molecules attached to loops formed by the mRNA. Each loop contributes to the signal by a constant *I*_0_. This constant is unknown and can vary from trace to trace due to noise in the experimental setup and the variability in the locations of the nuclei in the embryo. All models are written for the renormalized signal *F*(*t*) = *I*(*t*)/*I*_0_.

Because the fluorescent signal is produced by discrete polymerases that travel down the gene, we divide the gene into chunks of 150 base pairs, a length that corresponds to the irreducible space occupied by a polymerase on the gene (Fig. 3 in the main text). The positions the polymerase can occupy on the gene are labeled by an index 1 ≤ *i* ≤ *r*. The number of MS2 loops that have been formed by a polymerase that has reached a given position depends only on the MS2 gene construct and we define a deterministic function *L*_*i*_ for the whole length of the gene that describes the number of MS2 loops that have been produced by a polymerase at position *i*. In practice the exact number of loops is not an integer and varies from base pair to base pair so we take *L*_*i*_ as the average number of loops at this polymerase position (see Fig. 3 in the main text).

When the gene is fully loaded with polymerases (the number of polymerases is equal to the length of the gene divided by 150 bp), the fluorescence intensity is 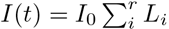. Assuming that the maximum of the signal over the whole trace is a good approximation for the fully loaded value we can determine *I*_0_ and renormalize the data. In practice, since we see variability in the expressed signal in different nuclei at the same position, we are not sure the fully loaded polymerase scenario occurs in each nuclei, so we take the mean of the maximum intensity values in the anterior. We use this renormalized fluorescence signal to infer the parameters of the dynamics.

The experimental data is analyzed assuming the system is in steady state and does not take into account the initial activation period after mitosis. and the end of the trace when the gene is deactivated before mitosis. We take only the middle window of the trace as shown in SIFig. 9.

In all models based on a stochastic gene switching (so all models except the Poisson model) we assume that the gene can be in several states with only two effective transcription rates: a non zero transcription rate in the ON state and an basal production rate equal to zero in the OFF state. When the gene is ON the polymerase loads at a maximal rate set by clearing of the binding site by the previous polymerase, which is one polymerase every 6 seconds (calculated as the irreducible polymerase length along the gene 150 bp divided by the polymerase speed, *v* = 25*bp/s*). The state of the gene is described by a stochastic process *X*(*t*) that is equal to 1 when the gene loads polymerase (i.e is ON) and 0 when the gene is OFF (see Fig. 1B in the main text). Once the polymerase is loaded its path is assumed to be deterministic with constant speed.

The gene can be described by the locations where there is a polymerase: we define *a*(*i*, *t*) as a function of time *t* and position 1 ≤ *i* ≤ *r* that is equal to 1 if there is polymerase at position *i* at time *t* and 0 otherwise (see Fig. 1D in the main text). The fluorescence signal is then a convolution of the polymerase position, *a*(*i*, *t*), and the details of the loop design of the MS2 construct, *L*_*i*_:

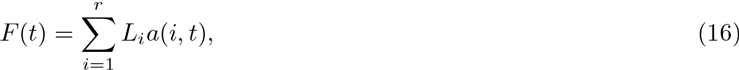

and the polymerase position can easily be translated back to the gene state through the deterministic relation, *a*(*i*, *t*) = *X*(*t* − *i*) (see Fig. 3D in the main text for the form of *L*_*i*_). This disruption is exact for a system with a discrete regulatory process and a discrete time step equal to the polymerase time step. Unfortunately, the moments in time when the gene switches are not necessarily multiples of the natural coarse graining steps of the system (the polymerase time step and its equivalent length) so it is necessary to introduce a continuous time in the system. We will present results for both the discrete and continuous time models. The continuous description is valid in the limit where the typical time spend by the gene in each state is long compared to the polymerase step or equivalently the gene switching constants are small compared to 1/6 s^-1^. See SI Section VIB for a more detailed argument.

**TABLE I:** The inferred *k*_on_ rates from the autocorrelation approach assuming a two state model for the four embryos and cell cycle 12 and 13, in the anterior and boundary.

| $k_{\text{on}}(1/s)$ | mov1 | mov2 | mov3 | mov4 |
| --- | --- | --- | --- | --- |
| 12A | 0.078 | 0.056 | 0.009 | 0.023 |
| 12B | 0.004 | 0.005 | 0.003 | 0.011 |
| 13A | 0.017 | 0.020 | 0.014 | 0.021 |
| 13B | 0.004 | 0.006 | 0.004 | 0.005 |

**FIG. 8:**
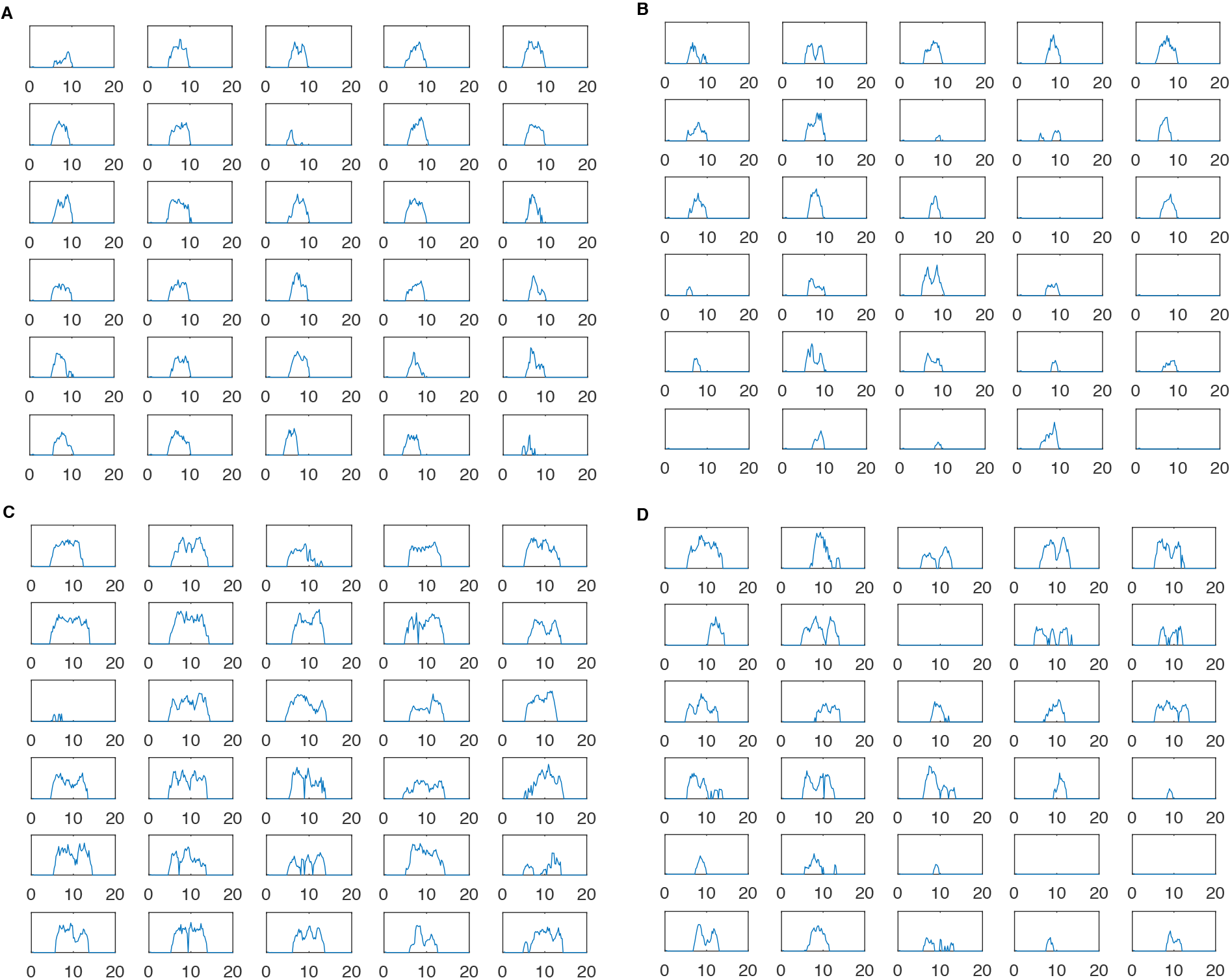
Examples of individual spot intensity over time. Consecutively shown are the traces in (A) Cycle 12, Anterior, (B) Cycle 12, Boundary (C) Cycle 13, Anterior, (D) Cycle 13, Boundary. The x axis is time in minute and y axis is the spot intensity in AU.

### B. The two state model

In this section we derive the equations required for the inference of the dynamics under the assumption that the gene can be in two states: ON or OFF represented by a two dimensional vector *x*(*t*) = [*x*_on_(*t*),*x*_off_ (*t*)]. *x*_on_(*t*) is the probability of the gene to be ON and *x*_off_ (*t*) is the probability for the gene to be OFF. *x*_on_(*t*) is the average over traces of the random variable *X*(*t*) depicted in Fig. 1B of the main text. We assume that the switching times between the two are exponentially distributed:

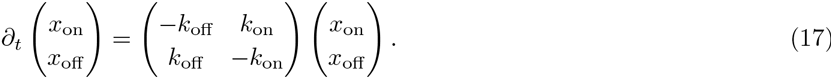

**FIG. 9:**
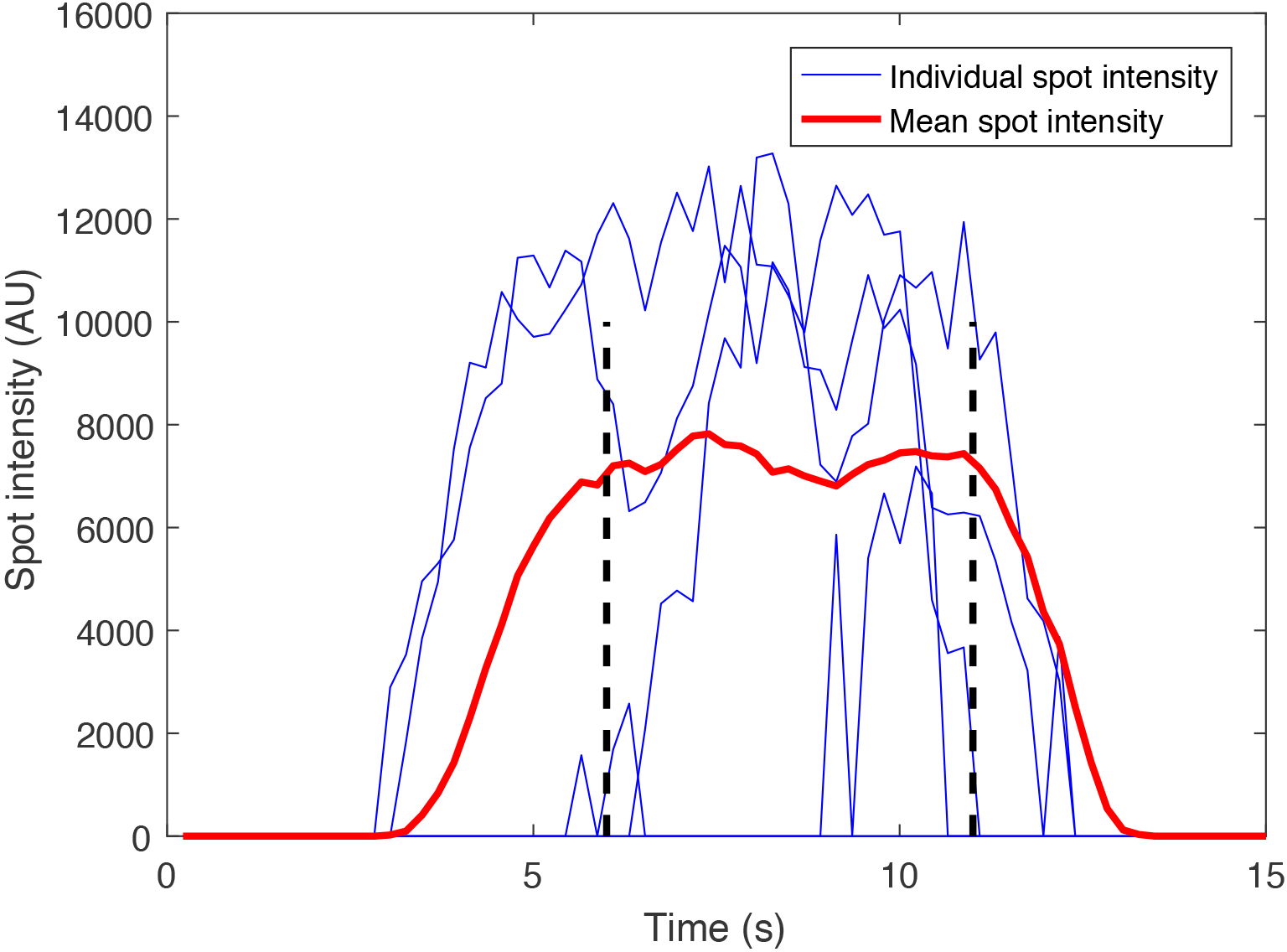
Data calibration. Shown are examples of 5 (out of 154) individual traces (blue) taken from embryo 1, cycle 13. Also shown is the mean spot intensity over time of all traces (red). The steady state window is chosen to be from the 6^*th*^ minute to the 11^*th*^ minute (dashed lines).

**TABLE II:** The inferred *k*_off_ rates from the autocorrelation approach assuming a two state model for the four embryos and cell cycle 12 and 13, in the anterior and boundary.

| koff(1/s) | mov1 | mov2 | mov3 | mov4 |
| --- | --- | --- | --- | --- |
| 12A | 0.060 | 0.088 | 0.008 | 0.019 |
| 12B | 0.020 | 0.034 | 0.021 | 0.051 |
| 13A | 0.018 | 0.031 | 0.016 | 0.027 |
| 13B | 0.031 | 0.054 | 0.031 | 0.064 |

The steady state probability to be ON is *P*_on_ = *x*_on_(*t* = ∞) = 1/*T*∑_*t*_ *x*_on_(*t*), where *T* is the duration of the stead state window in Fig. 9, and is:

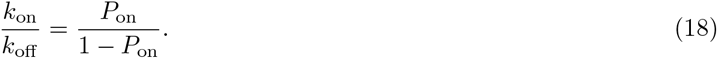

We learn *P*_on_ from Eq. 1 in the main text:

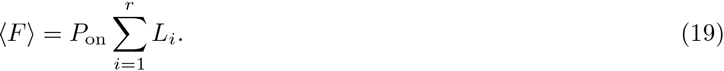

and use it to obtain the ratio of the switching rate from Eq. 18.

**FIG. 10:**
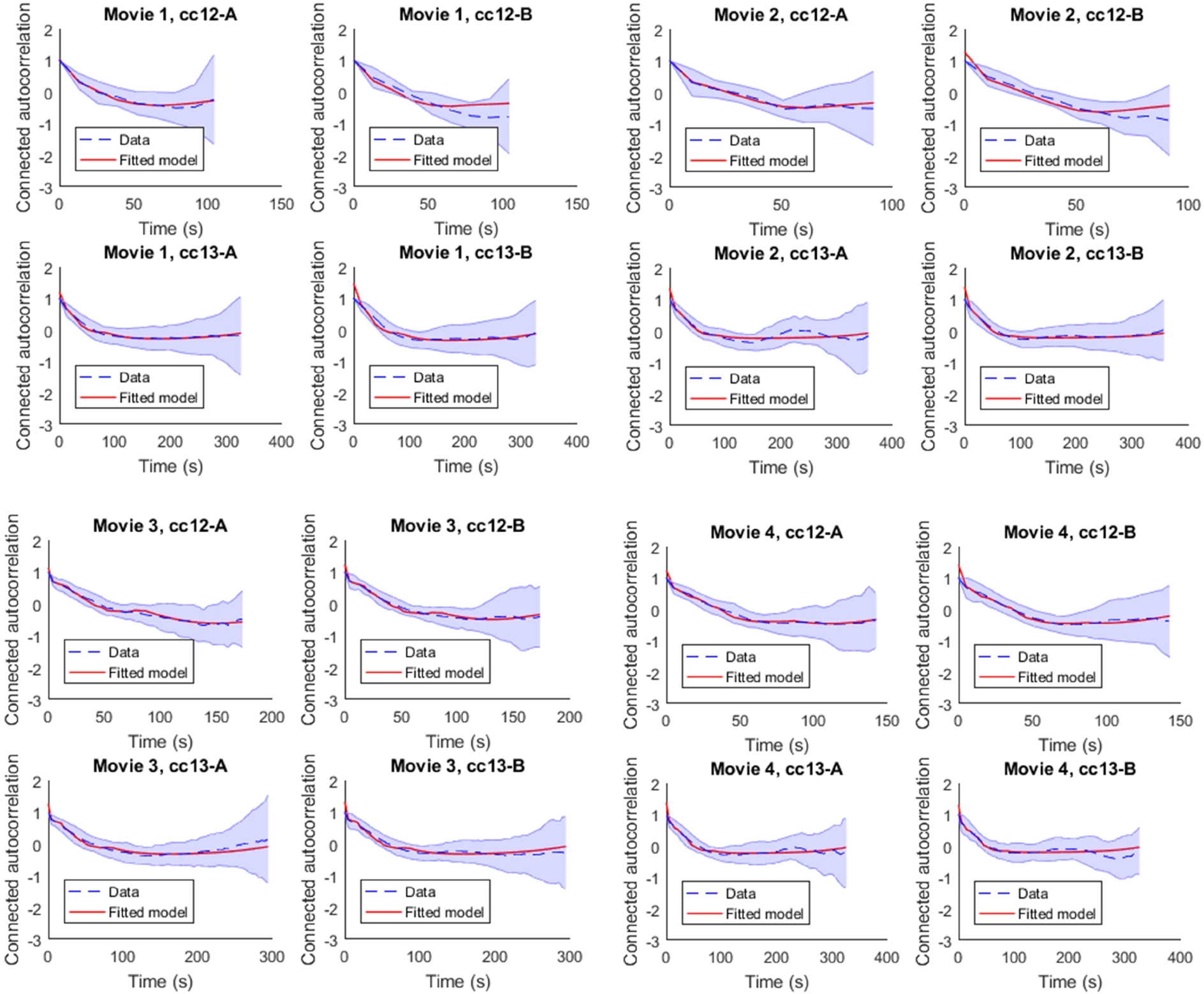
Fits of the autocorrelation function. The empirical autocorrelation function for both the anterior and boundary regions in all four embryos is fit using the autocorrelation function with the finite size corrections for the two state model.

The autocorrelation function is:

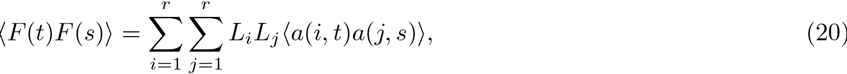

where the brackets are an average over traces (different realizations of the random process). We define *A*(*t* – *i*,*s* – *j*) = 1/*x*_on_(*s* – *j*)〈*a*(*i*,*t*)*a*(*j*,*s*)〉 – the probability that the polymerase is at position *i* and time *t* given that there was a polymerase at position *j* at time *s* (here we assume that *t* – *i* ≥ *s – j*). Using the deterministic relation between the polymerase position at a given time *a*(*i*, *t*) and the probability to be on at an earlier time *X*(*t* − *i*), *A*(*t* − *i*, *s* − *j*) is equivalent to the probability that the gene in ON at time *t* − *i* given that is was ON at time *s* − *j*:

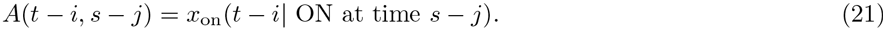

Plugging the expression into Eq. 20 we obtain Eq. 2 in the main text:

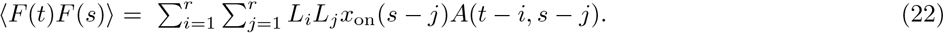

In steady state the system is translationally invariant *A*(*t* − *i,s* − *j*) = *A*(|*t* − *i* − *s* − *j*|) and for brevity we will denote is as *A*(*n*) – the probability that the gene is ON at time *n*, given that it was ON at time 0. To find *A*_*n*_ we need to solve for *x*(*t*):

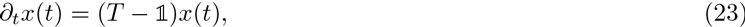

where 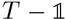 is given by Eq. 17 and calculate the expectation value that the gene is ON at time *t* given in was ON initially:

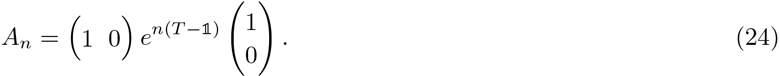

**FIG. 11:**
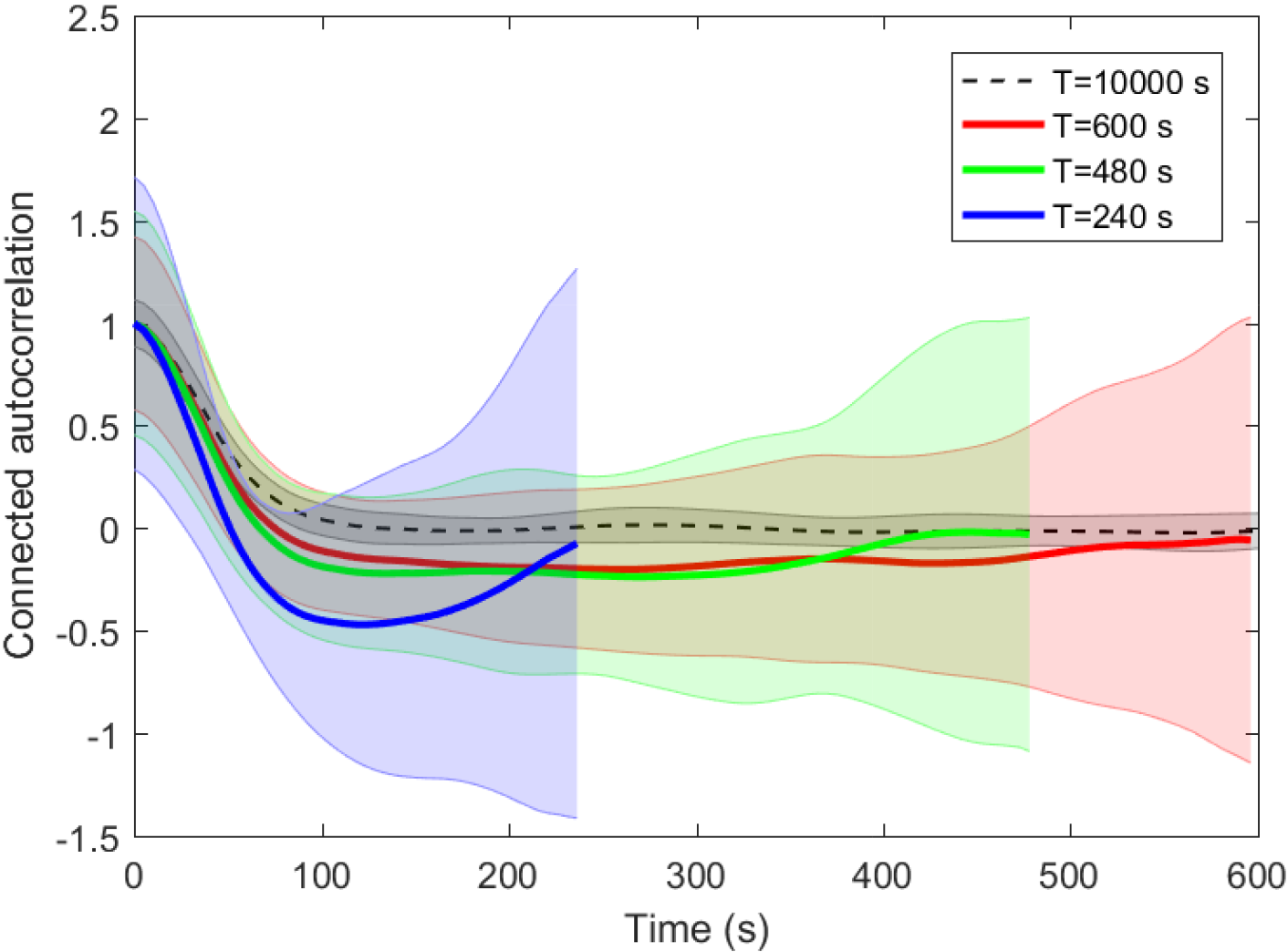
Example of the connected autocorrelation function for the two state model calculated for different trace lengths as a function of time *T*. The shaded areas denote the standard variation over *xx* simulated traces. The switching rates *k*_on_ = *k*_off_ = 0.01*s*^-1^ and the number of nuclei *M* = 500.

Eq. 24 is correct in a continuous time model. Its discrete time equivalent is

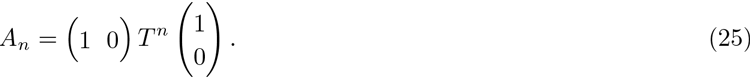

In the limit of *k*_on_ and *k*_off_ much smaller than the polymerase step they are also much smaller than 1 and 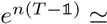 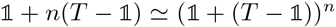. In this limit the continuous and discrete time descriptions of Eq. 24 and Eq. 25 are equal.

The eigenvalues of 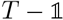 are [1, *δ*], where *δ* = 1 – *k*_on_ – *k*_off_ with corresponding eigenfunctions:

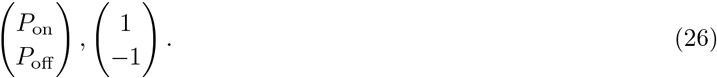

The transition matrix *T* is

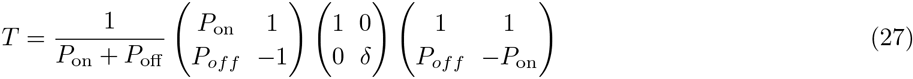

and

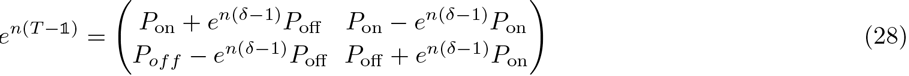

resulting in

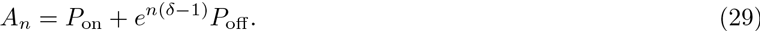

In steady state *x*_on_(*s – j*) = *P*_on_ and the connected autocorrelation is:

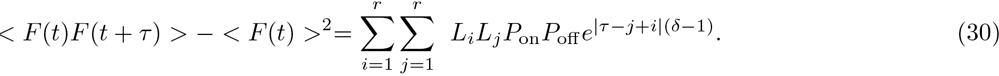

Since we already know the ratio of the rates from *P*_on_, inferring *δ* using Eq. 72 determines *k*_on_ and *k*_off_.

### C. Computing out of steady state

The autocorrelation approach can be generalized to a case when the system is out of steady state, when the autocorrelation function explicitly depends on the two time points and not only on their difference. During mitosis the gene is OFF and then gets turned ON in early interphase. Motivated by the hunchback expression we will present the calculation assuming the gene is initially ON, but it is generalizable to any other initial condition. Assuming *t – i* < *s – j*, we want to calculate the probability that the polymerase is at position *i* at time *t*, given that it was at position *j* at time *s*. Since the gene is initially OFF, we need to calculate the probability that the gene is ON at time *t – i*. The autocorrelation function of the polymerase position is:

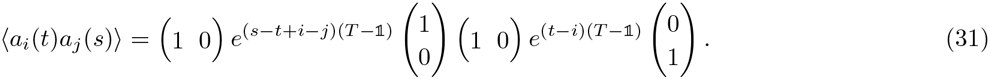

Using Eq. 29 and

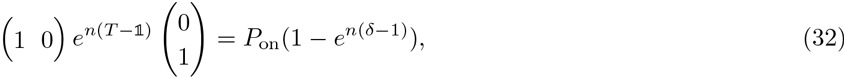

we obtain:

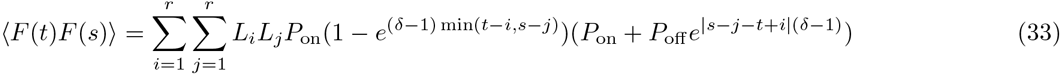

### D. Multiple off states

The calculations presented in Appendix VIB can be extended to models that include more OFF or ON states as long there are only two production states for the mRNA: one enhanced and one basal production state. The transition matrix *T* will then be of higher dimension and in practice should be (and has to be for dimensions larger than 3) diagonalized numerically. The exact analytical solution for the autocorrelation function is still valid written in terms of the powers of *T*.

### E. Generalized multi step model

A gene with many OFF states can also be described using a reduced model with two effective gene expression states ON and OFF, where the times of transitions between these two state are not exponential but follow a long tailed distribution approximated by a Gamma distribution. The Gamma distribution describes an effective transition over many irreversible transitions between a series of OFF states:

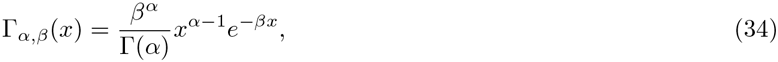

where *β* is the scale parameter, *α* is the shape parameter, and Γ(*α*) is the gamma function. The mean time time spent in the OFF state is 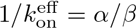, so the probability for the gene to be in the ON state is:

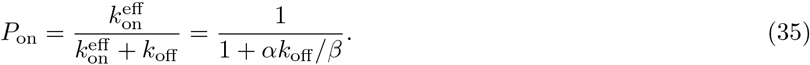

This model has three parameters, regardless of the number of OFF states, and using Eq. ?? reduces the number of parameters to two, which greatly simplifies the inference. The remaining two parameters are learned from the autocorrelation function in Eq. 20, which formally has the same form as Eq. 22:

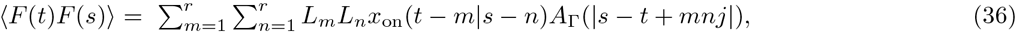

but *A*_Γ_(|*s – t* + *m – n*|) = *x*_on_(*t – m*|*s – n*) is now not memoryless. We limit our presentation to the steady state, but the calculation generalizes to out of steady systems.

We cannot solve the problem in real space, but we compute the Fourier transform of the autocorrelation function of the fluorescence signal:

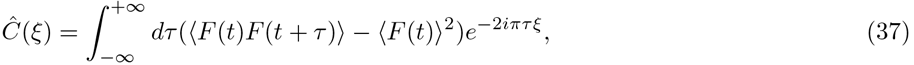

which using Eq. 20

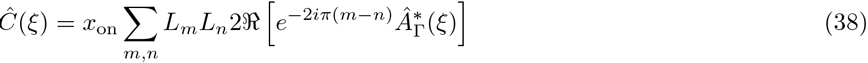

we reduce to calculating

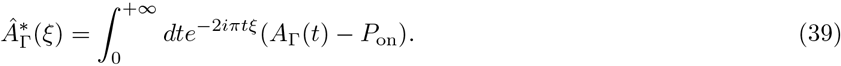

We decompose *A*_Γ_(*t*) into a sum over full cycles of the gene turning from ON to OFF, with the constraint that at time *t* the gene is ON:

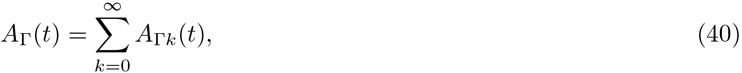

where

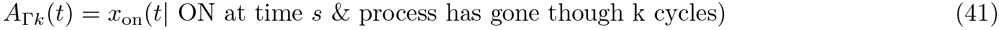

Since the first jump is from the ON to OFF, which is exponential it contributes 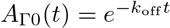.

First we compute an auxiliary probability distribution function of the time it takes the process to go through a full ON-OFF cycle η(*t*) of taking an exponential jump out of the ON state followed by a Gamma distributed jump out of the OFF state:

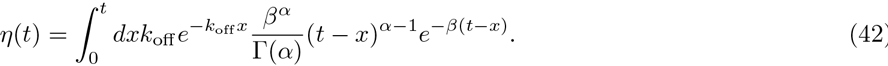

The Fourier transform of this distribution is:

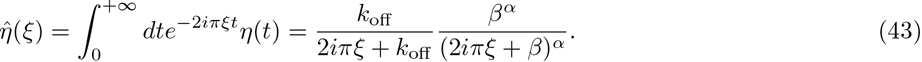

To compute 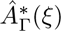 we need to sum over all the possible times at which the cycles could have occurred, with the constraint that at time *t* the gene is ON:

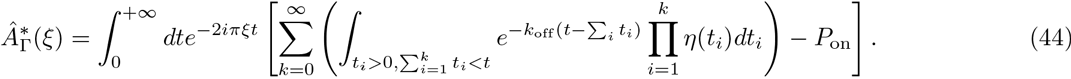

We can rewrite the last term in Eq. 44:

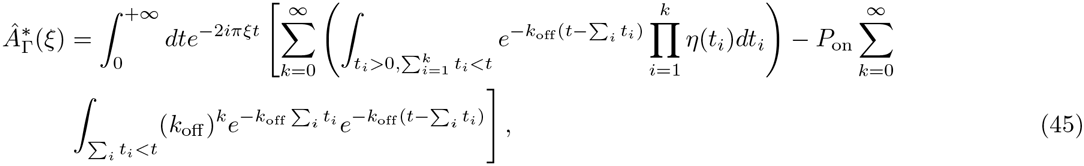

using the expansion of unity:

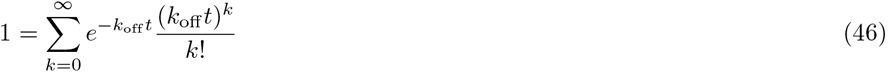

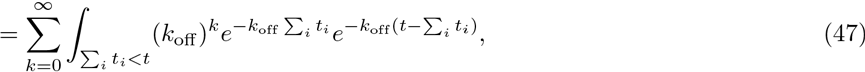

with the convention for the *k* = 0 term:

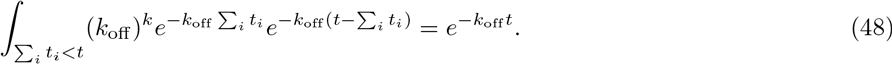

Collecting terms:

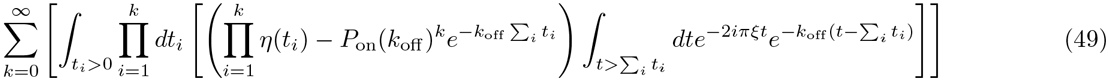

and setting *u* = *t* ∑_*i*_ *t_i_* in the last integral:

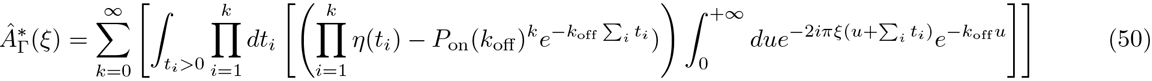

we obtain:

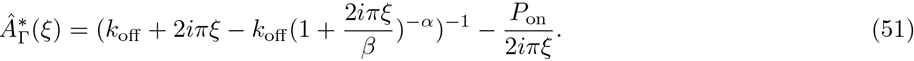

Using Eq. 36 we recover Eq. 11 in Materials and Methods. For *α* = 1 we recover results of the two state model.

### F. The autocorrelation of a Poisson polymerase firing model

We compared the auto-correlation function for our models with bursty dynamics to the auto-correlation of a model that assumes in steady state stochastic gene expression with a constant exponentially distributed rate – a Poisson polymerase firing model. We assume that the gene expression rate is memoryless and the transcription interval follows an exponential distribution of mean τ_*P*_:

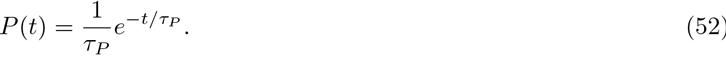

In order to compare the two models we need to reinterpret the statistics introduced for bursty dynamics in the framework of a Poisson model. The quantity *P*_on_ corresponds to the average occupancy of polymerase sites on the gene. This constant can be computed for a Poisson arrival model. The size of the polymerase is 150 bp and its speed is ~ 25 bp/second, the maximum loading rate of polymerase is one every 6 second. Since the polymerase cannot load faster than once every 6 seconds, we calculate the average occupancy of the gene as the temporal average of probability that the polymerase starts transcribing within 6 seconds:

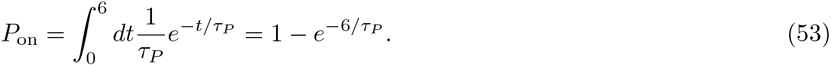

Here we assume that the next polymerase to bind can be recruited while the previous one is clearing off the binding site. Since the process is memoryless and the Poisson firing process is uncorrelated, its connected autocorrelation is close to a delta function δ(τ = 0). However, due to the gene lengthy elongation time, there is a non-flat autocorrelation function of the fluorescence signal. the probability of the polymerase to be at position *i* at time *t*, given it the gene to ON as predicted from the MS2 signal at short times. At steady state, the connected auto-correlation function is:

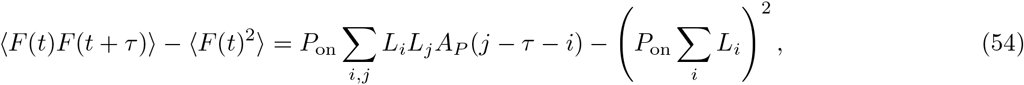

where *A_P_*(*τ*) is the probability of the polymerase to be at position *i* at time *τ*, given it was at position *j* at time 0 in the Poisson firing model.

If τ < 6*s* then the two positions on the gene, *i* and *j*, share the same polymerase with a probability proportional to |6 – τ|, taking equally distributed polymerase positions. If τ > 6*s*, *A_P_*(*τ*) is given by the probability that there is a polymerase at the second site, which is independent of what happened at the first site. The two cases give:

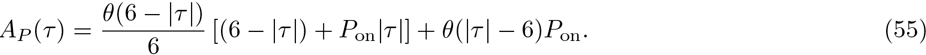

This function is flat for τ > 6*s* and the first part of the right hand side of Eq. 55 has little effect on the autocorrelation function over a cell cycle (as cell cycle duration is much bigger than 6*s*). For this reason we use a flat function as a very good approximation for *A_P_* in our analysis.

From the form of Eqs. 54 and 55 and the flat approximation of *A_P_* we see that *P*_on_ is only a normalizing constant and the shape of the function is completely determined by the loop function *L*_*i*_, which is known. We can compare the expected autocorrelation function of a Poisson model to data and find that it does not explain experimental results as well as bursty dynamics (although gene switching models have higher numbers of parameters).

From Eq. 53 we can learn polymerase arrival rates in the anterior and at the boundary of the embryo. We find that the Poisson model would require very high heterogeneity of polymerase arrival times as a function of A-P axis position. At the boundary in particular we expect the mean polymerase arrival time to be above 60*s*.

### G. Numerical simulations

To simulate the time evolution of MCP-GFP loci's intensity, we used the Gillespie algorithm [41, 42] to predict the time it takes for the gene to switch between the states, the active ON state and the inactive OFF states. In all models we assume that the time of the transition from the active to the inactive states, τ_on_ is exponentially distributed with rate *k*_off_. The time of the transition from the inactive OFF states to ON state, τ_off_ depends on the model considered:

- For the two-state model τ_off_ is exponentially distributed with rate *k*_on_.
- For the three-state model τ_off_ is a sum of two exponential processes with rates *k*_1_ and *k*_2_ that describe the transitions between the two OFF states.
- For the Gamma model τ_off_ is chosen from a the Γ(*α*, *β*) distribution defined in Eq. 34.

To generate the traces of length *T* from *N* nuclei, we first simulate a long trajectory of length *N × T*, denoted as *X*(*t*). To account for the incompressibility of the polymerase, we divide the traces into 6s intervals, which is the time the polymerase needs to cover a region of the gene equal to its own lengths. We assume that at each 6s time point, if the gene is in the ON state, there is a transcription initiation event by a single RNA polymerase with a full transcription rate, defined as the length of the gene divided by the polymerase velocity, defined in SI section VI A. Following this event, the RNA polymerase will slide along the target gene segment and synthesize a nascent RNA. At time *i* into this elongation process, the nascent RNA has *L*_*i*_ MS2 binding sites as depicted in Fig. 3 of the main text. To impose 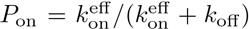 If the gene switches into the OFF state before a full 6s interval, the polymerase transcribes the gene at a reduced rate proportional to the fraction of the 6s interval for which the gene was ON. The number of MS2 binding sites at the transcription locus site is therefore given by the convolution of the gene state and the promoter construct design function *L* (see Fig. 1 in the main text):

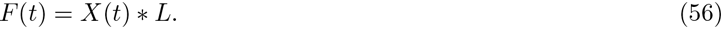

We assume that the number of MCP-GFP molecules in the nuclei is sufficient to bind to all newly transcribed MS2 binding sites and that the binding process is infinitely fast. The spot intensity is calculated as the number of binding sites produced at the loci (given the intensity of each MPC-GFP dimer equal to 1). Lastly, the long spot intensity traces are divided equally into *N* smaller traces of length *T*.

### H. Correction to the autocorrelation function for finite trace lengths

The short duration of the experimental traces, *v*_*α*, *i*_, where 1 ≤ α ≤ *M* describes the identity of the trace and 0 < *i* < *K* denotes the sampling times, coupled with the need to correct for experimental biases by calculating the connected correlation function introduces finite size effects. The true connected correlation function between time points at a distance *r*, *C*_*r*_ (red line in Fig. 12), is not equal to the empirical connected correlation function calculated as an average over the *M* traces, *c*(*r*) (blue line in Fig. 12), of the autocorrelation functions of the finite traces. The theoretical connected autocorrelation function calculated in our model is:

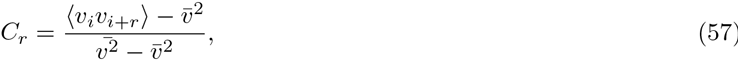

where 〈·〉 denotes an average over random realizations of the process and we assume steady state 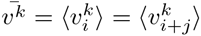.

The empirical connected correlation function of each finite trace of length *K* << ∞ has the form:

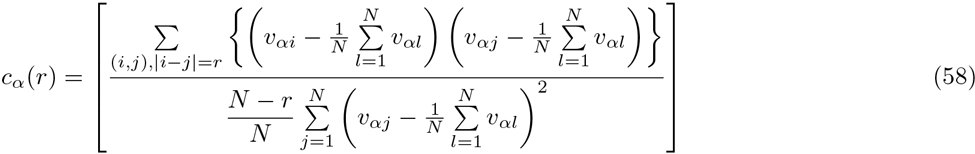

and the empirical connected correlation function calculated averaged over *M* traces is

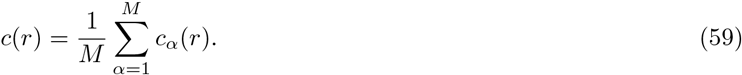

*C*_*r*_ requires knowing the true second moment of the fluorescence signal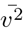. In our data we find that the true variance of the normalized fluorescence signal, 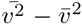 is well approximated by the average over traces, so we approximate Eq. 58 by:

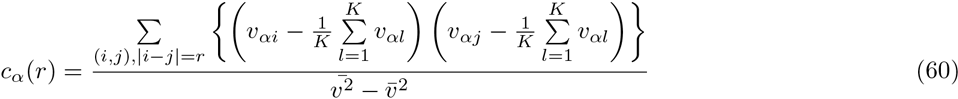

The difference between the theoretical and empirical connected correlation function is independent of our model and arises for the connected correlation function of any random process, as shown in Fig. 12 for the simplest random process – the Ornstein-Uhlenbeck process. The difference is due to the fact that the short time average induces spurious correlations when calculating averages of the signal taken at different times. When analyzing the data, to avoid describing nucleus-to-nucleus variability that is not connected to the signal, we first subtract the mean steady state fluorescence signal of each trace, normalize this connected autocorrelation function to 1 at time *t* = 0, and then average over traces (Eq. 60) before averaging over the trace ensemble (Eq. 59). In steady state, the infinite trace mean equals the ensemble average, 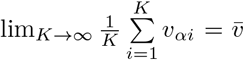. However, as shown in Fig. 2 of the main text, the short trace mean is not a good approximation to the long term (or ensemble) average, 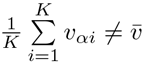. The points located in the center of the trace are much more correlated with the mean than the points at the beginning and end of the time interval. The correction for each value of *r* is different and must be separately computed.

In analyzing our data we use the finite size correction for the mean derived below that expresses the empirical connected correlation function *c*(*r*) in terms of the theoretical connected correlation function *C*_*r*_. For *K* → ∞ the empirical connected correlation function becomes the infinite time connected correlation function, however our traces are very short. These corrections are valid for all time dependent data sets so for completeness the finite size correction for the variance is derived in SI Section VI I but is not used in the analysis.

The number of pairs of time points of distance *r* in a trace of length *N* is simply *N* − *r* and the combination of Eqs. 59 and Eqs. 60 becomes:

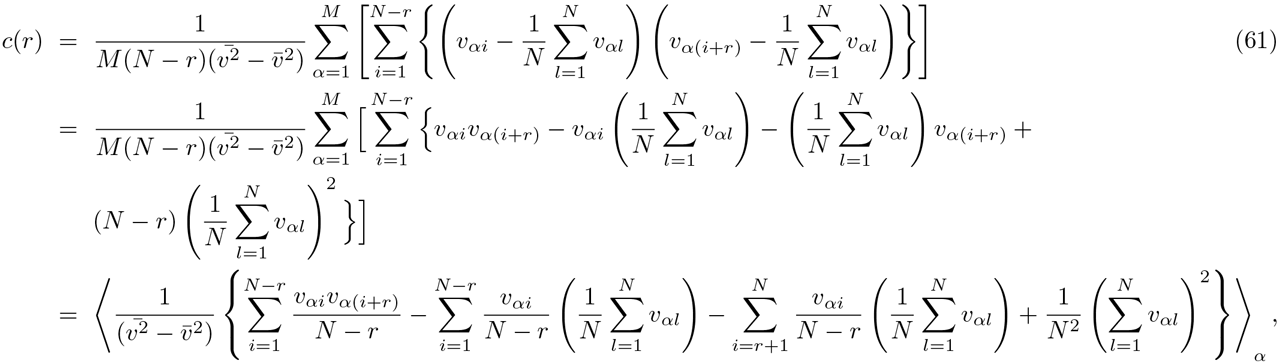

where we have explicitly written out the terms and in the last line we introduced the average over traces 〈·〉_α_ = 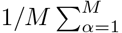. In steady state due to time invariance:

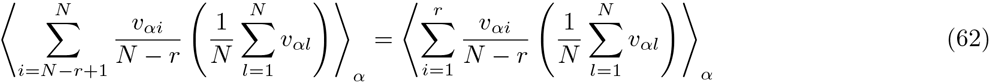

and the theoretical (not connected) correlation between two points is a function only of the distance between these two points:

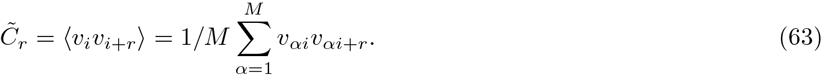

We have assumed that *M* is large and a population average over the *M* traces for points separated by *r* on each trace approximates the *M* → ∞ limit of the theoretical average over different realizations of the process. Using Eq. 63 we obtain:

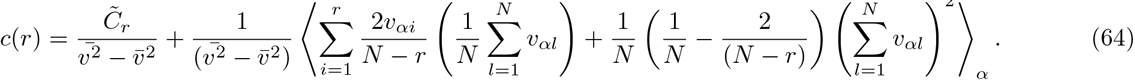

To rewrite 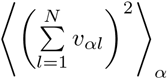 as a sum over *C*_*r*_ we calculate the number of pairs of time points separated by a distance *k* in the whole trace of length *N*. For *k* = 0 it is equal to *N* and for 1 ≤ *k* ≤ *N* – 1 it is equal to 2(*N – k*):

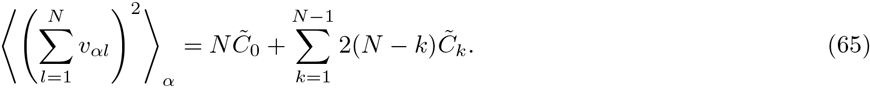

Similarly

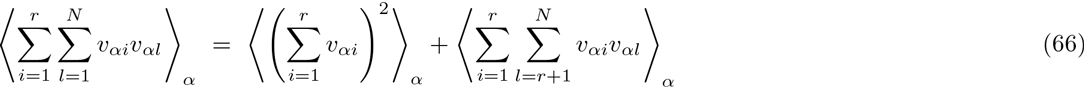

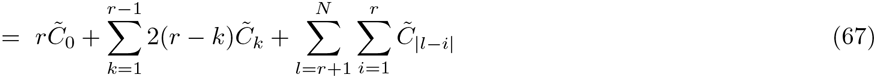

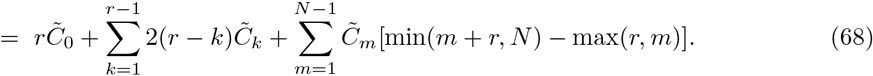

**FIG. 12:**
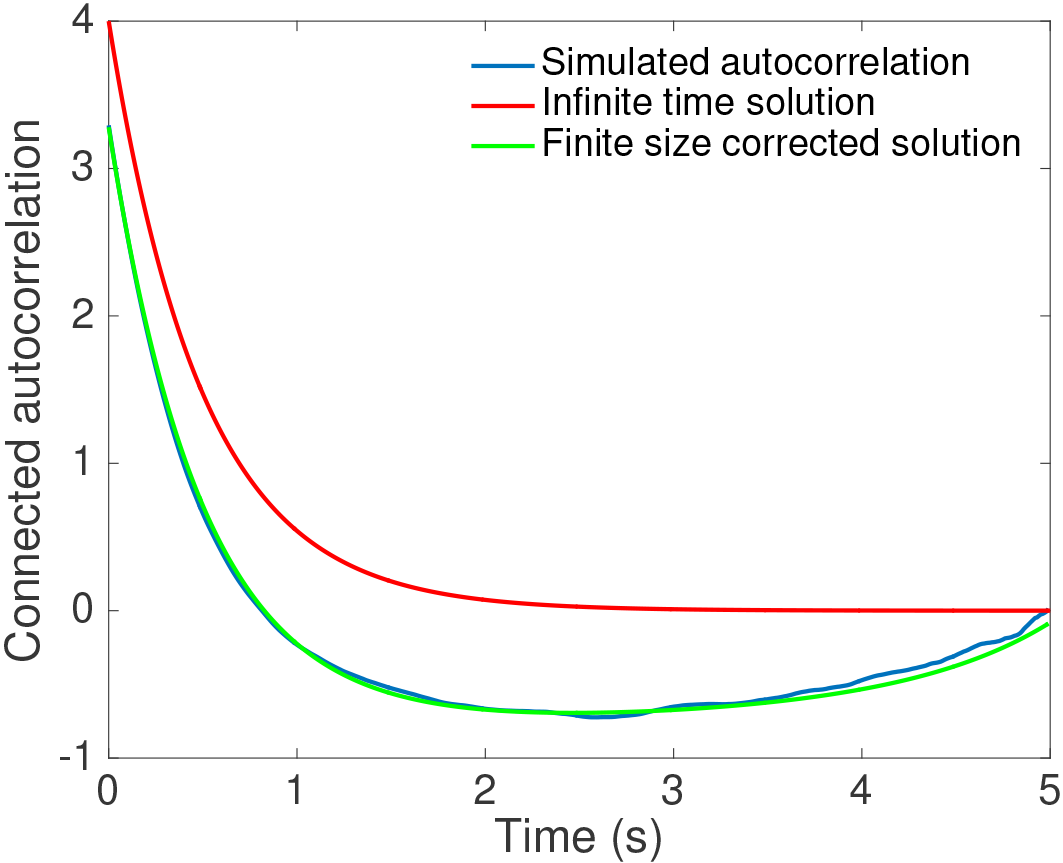
The finite trace effect for the Ornstein-Uhlenbeck process. The connected autocorrelation function *C*_*r*_ = exp(-*t*/τ) (red line) compared to the connected autocorrelation function calculated from short time traces as described in SI Section VIH (blue line) and the corrected connected autocorrelation function (Eq. 69 green line). λ = 2s^-1^, γ = 4s^-1/2^ and the short trace length is 5s where the Ornstein-Ulhenbeck process is ∂_t_*x* = −λ*x* + γξ and ξ is Gaussian white noise.

Collecting the empirical connected autocorrelation function in Eq. 59 is expressed in terms of the theoretical nonconnected correlation function in Eq 63 as:

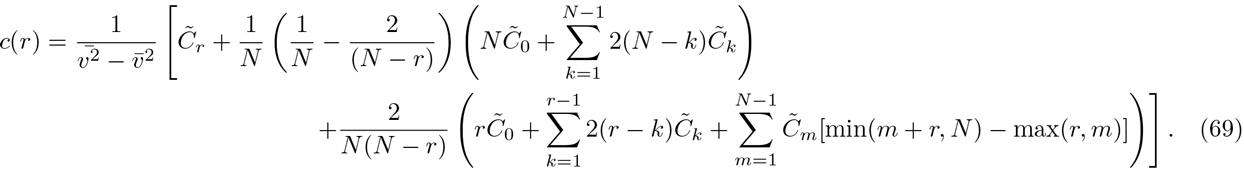

### I. Correction to the autocorrelation function from correlations in the variance

In SI Section VI H we calculated the finite size correction due to short traces for the empirical connected correlation function assuming that differences between he empirical variance and the theoretical variance for infinite traces do not affect the connected autocorrelation function. This approximation is valid for our data. For completeness we now calculate the finite size correction coming from spurious correlations in the variance obtained when computing the variance trace by trace, before averaging over the traces (Eq. 59). Analyzing the data, we normalize the autocorrelation function of each trace before taking the average over all traces because of potential nucleus-to-nucleus variability in the signal calibration. This is equivalent to dividing each autocorrelation function by its variance, before averaging over the traces and can introduce errors.

The empirical connected correlation function in Eqs. 59 and 58 can be rewritten by adding and subtracting 1 in the denominator as:

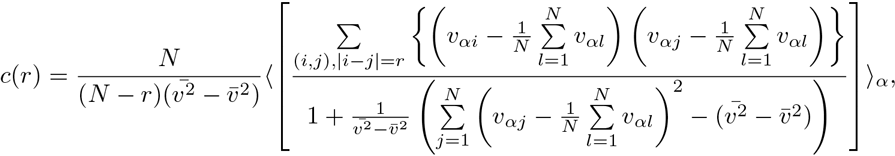

where the average 〈·〉_*α*_ is over *M* traces as defined in SI Section VIH. Assuming the true variance of the process is close to the empirical variance we linearize the denominator:

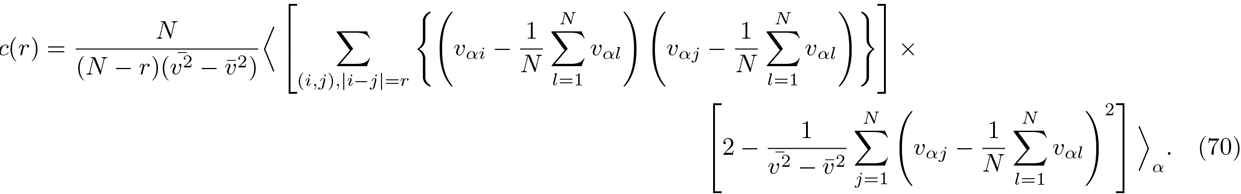

We first term is proportional to the connected correlation function in Eq. 69 we calculated in SI Section VIH assuming constant variance. We focus on the second term:

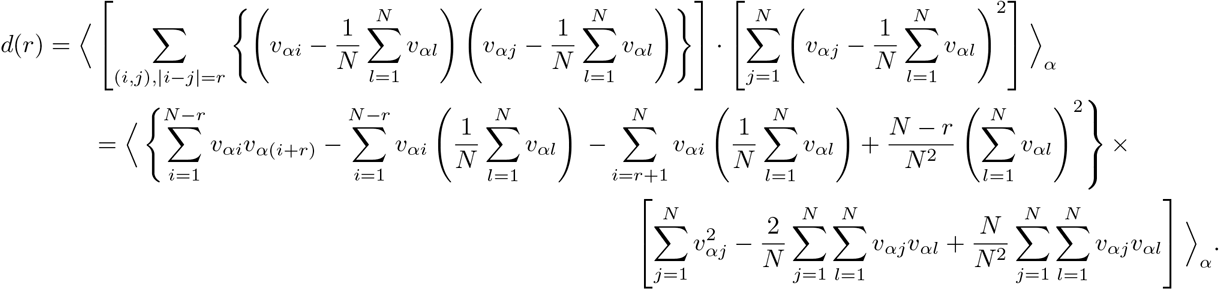

Using time invariance at steady state (Eq. 62) in the first factor and simplifying the algebra in the second factor:

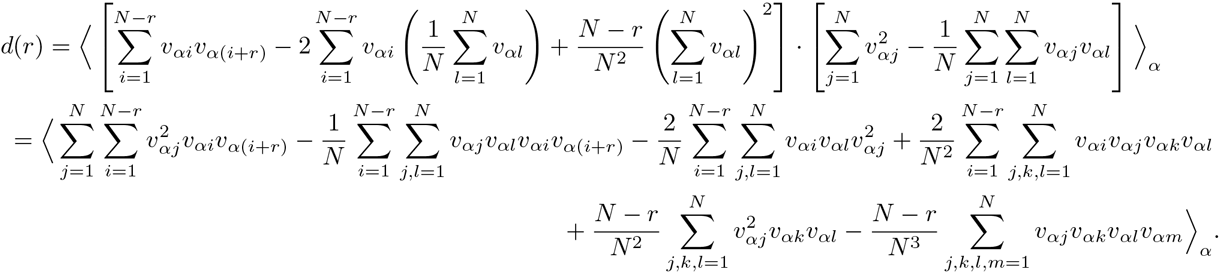

The final correction for correlation due to correlations in the variance coming from short time traces is easily evaluated terms of four-points correlation function 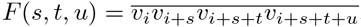.

### J. Cross-correlation

The presented correlation analysis can also be extended to constructs with two colored promoters inserted at two difference positions on the same gene. In this case, each construct can have a different loop design function 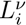, where *v* =1, 2, and the cross-correlation of the normalized fluorescence intensity is:

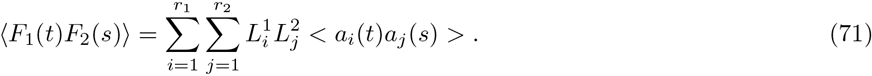

The 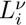 functions start at the same point (the one describing the downstream construct is 0 for the first steps).

After the loop design functions 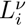 have been defined, the calculation of the theoretical cross-correlation function and auto-correlation rely only on calculating the correlations of the gene expression state, which is the same for both. So the results presented for the particular models are valid, after correcting for the two different loops functions. For examples, the steady state connected cross-correlation function of the two state model is:

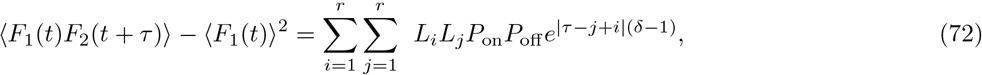

where *P*_on_ and 〈*F*_1_(*t*)〉^2^ = 〈*F*_2_(*t*)〉^2^ can be independently calculated from either probe, which provides an independent estimate of the experimental noise.

The differences in the use of the cross-correlation function and auto-correlation function arise when calculating the finite size corrections from short traces, because assumptions about the statistical time invariance of the signal in steady state are no longer valid. The non-connected theoretical correlation function (equivalent of Eq. 63) is now defined on two signals, *v_i_* and *w_i_*:

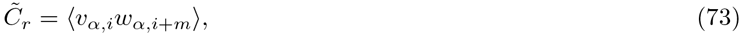

where 〈·〉 define the average over random realizations of the process and in steady state is independent of *i*. Unlike for the auto-correlation function, 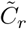 is no longer symmetric with exchange of *v_i_* and *w_i_*. The empirical cross-correlation function is (assuming the variance is well approximated by the empirical variance):

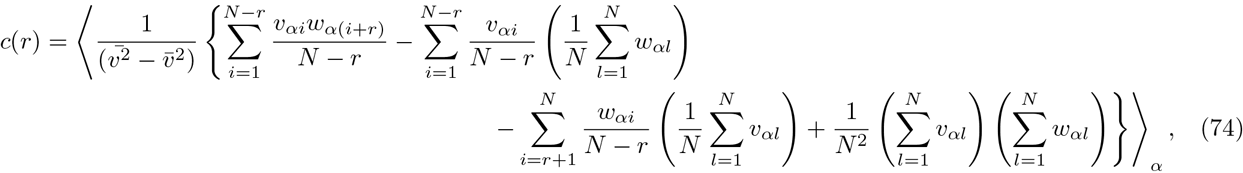

which in terms of the 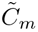 is:

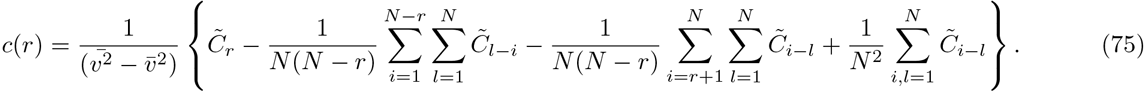

Repeating the steps in SI Section VIH we obtain the finite size correction for the cross-correlation function.

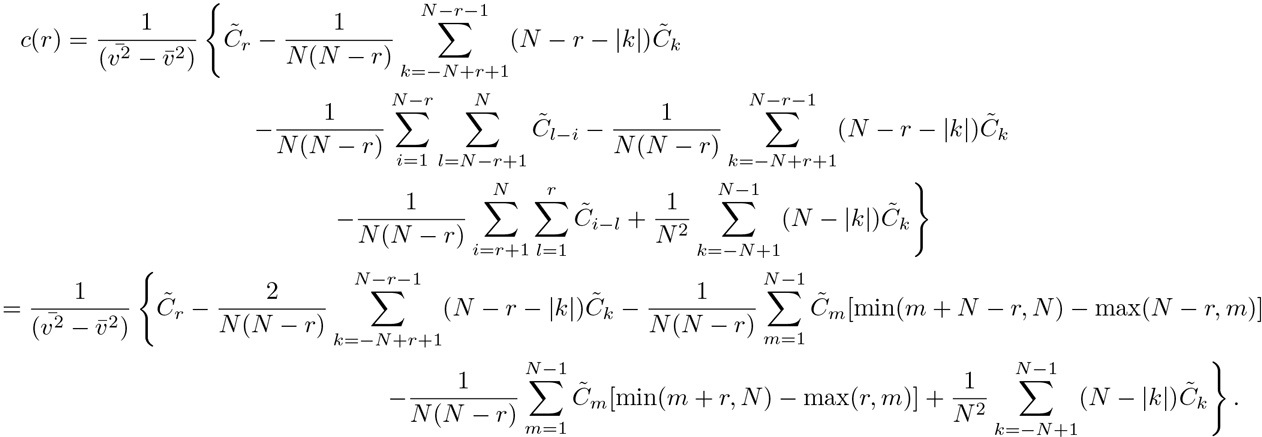

### K. Precision of the translational process

The precision of the total mRNA produced during a cell cycle presented in the main text is proportional to the activity of the gene and requires a careful calculation of the variability of the probability of the gene to be ON in different nuclei at the same position. The total activity of a nucleus, defined as the integral of the normalized fluoresce 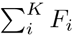, where *i* < *K* are the sampling times in steady state window of the cycle, in steady state is proportional to the probability of the gene to be ON in a given trace, 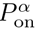. To keep our analysis independent of normalization, we will calculate the relative error defined as the variance over the mean of 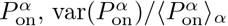, where the averages are taken over traces.

First, we can calculate the relative error of the probability of the gene to be ON 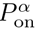 directly from the traces. We compute the mean and standard deviation of the distribution of 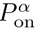 in a given window along the AP axis. 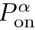 for each trace is calculated from Eq. 19.

**FIG. 13:**
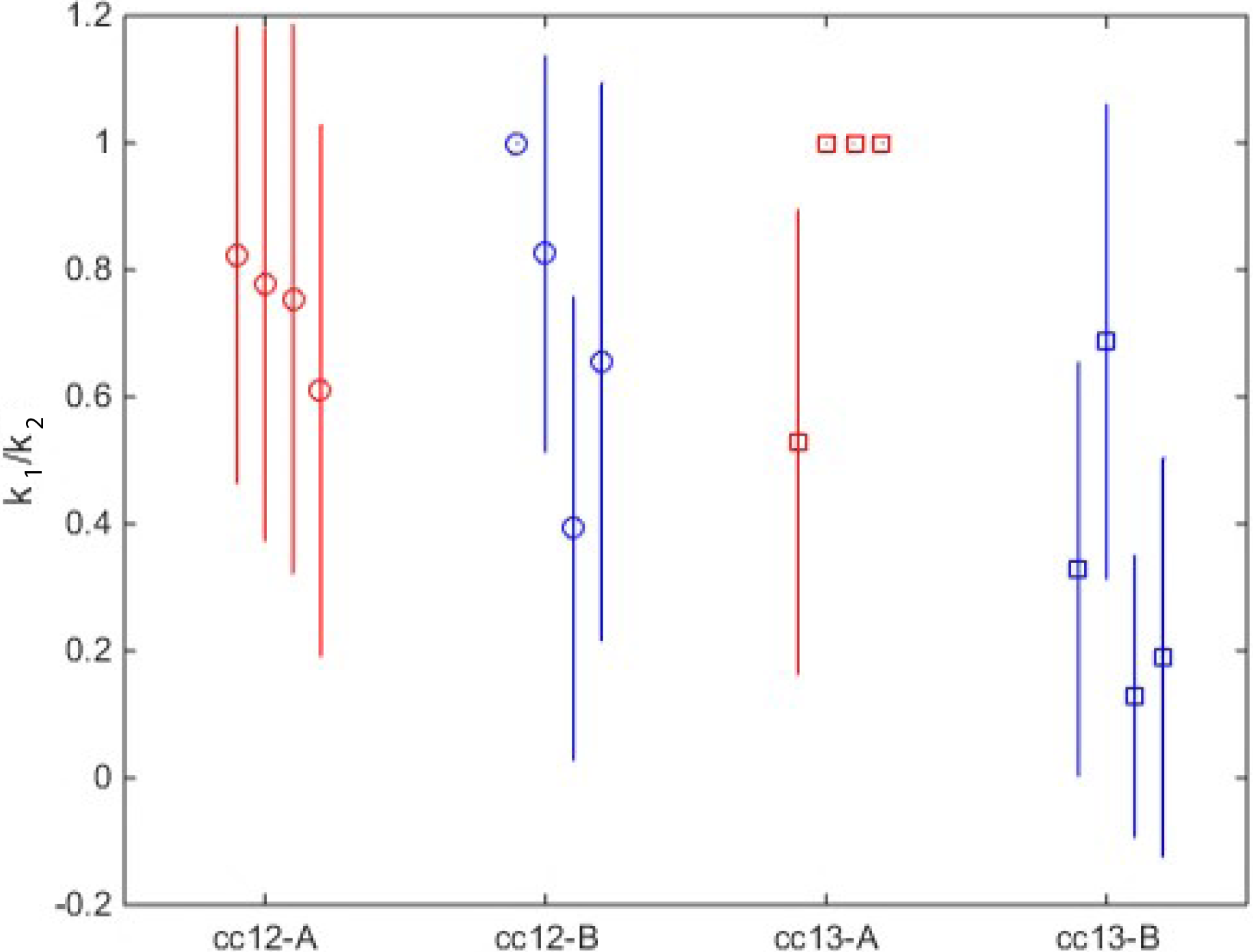
The fit of the three state cycle model to the data. The fit of the ratio of the two rates for leaving the two OFF states *k*_1_/*k*_2_ to the steady state traces from four embryos in the anterior and boundary region of cell cycle 12 and 13. Each point is data from one embryo. The error bar represent the standard deviation of the inferred value. The fit is for a randomized 60% of the data. The sum of the switching rates *k*_on_ + *k*_1_ + *k*_2_ is shown in Fig. 5B of the main text.

We can compare the results of the empirically estimated relative error to predictions of the steady state models. We know that the expected average over traces 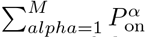 is *P*_on_. Within the assumption of our model presented in SI Section VIB, the expectation value of the square of the 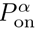 is expressed in terms of the expression states of the gene, *X*(*t*):

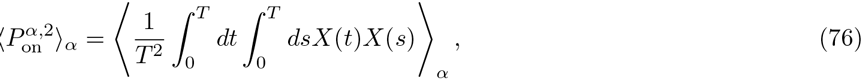

where the average is over *M* traces and *T* is the total duration of the trace in real time. In terms of the probability that the gene is ON at time τ given that it was ON at time 0, *A*(τ) defined in Eq. 21, we obtain

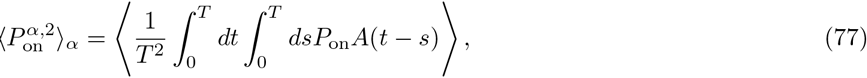

where *A*(τ) has units of seconds. The relative error is obtained by replacing *A*(τ) by the appropriate function for each model. For the two state model:

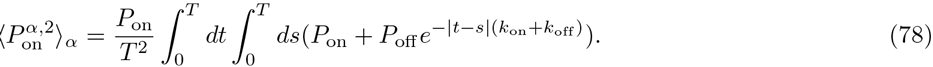

**FIG. 14:**
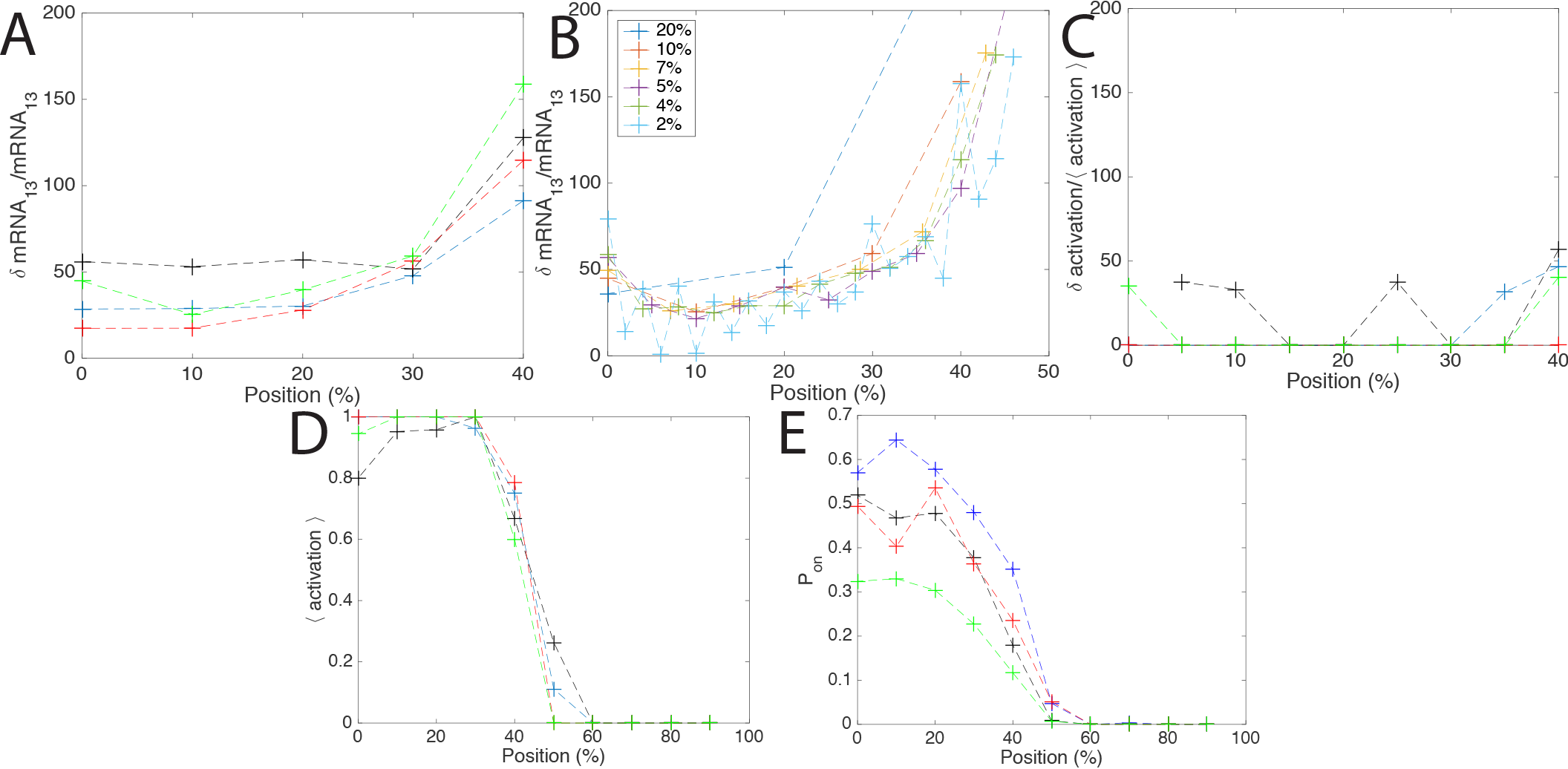
The relative error of gene expression. A. The conclusions about precision do not depend on the embryo. The relative error of the total mRNA produced in cell cycle 13 as a function of position for windows equal to 10% of the embyo length. Each colored line represents one embryo. The same data plotted as an average over embryos with the variance as error bars is shown in Fig. 7 of the main text. B. The conclusions about precision do not depend on the window size. The total mRNA produced in cell cycle 13 as a function of position for different window sizes. Except for very large scales (20%) and very small scales comparable to one nuclear width (2%, the relative error as a function of position is reproducible. C.The relative error of the discrete variable that describes the probability of the gene to be ON at any time during the cell cycle as function of position. The relative error is much lower in the anterior compared to the error in the total produced mRNA, but remains high at the boundary. D. The mean probability of the gene to be ON at any time during the cell cycle as a function of the embryo length (binary approximation). E. The mean probability for the gene to be ON averaged over the cell cycle. In C-E each colored lines describe different embryos.

Integrating and substracting the mean squared we obtain the relative error:

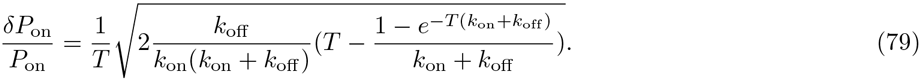

The probability of the gene to be on is proportional to the total mRNA produced and for large *T* we reproduce the result in Eq. 4 in the main text:

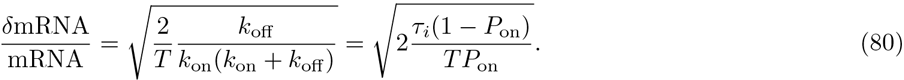

For the three state cycle model the same calculation is valid until Eq. 77 and is then carried out numerically.

Precision from static (Fluorescent In Situ Hybridization – FISH) images is calculated as the variance over the mean of the distribution of a binary variable, which for each nucleus is 1 if the gene is on in the static image and 0 if it off [5, 6, 24]. The signal in FISH datasets in an average over an unknown timeframe. To compare our analysis of the time dependent signal to these previous measurements, we use a binary variable, which is 1 for each nucleus that was ON during the steady state interphase and 0 for each nucleus that was always OFF. The results of the relative error as a function of position obtained using this empirical analysis in SIFig. 14 show agreement with previous reports [6]: for most traces the relative error in the anterior is zero – all nuclei in a given AP axis window express, and it increases to ~ 50% at the boundary.

**FIG. 15:**
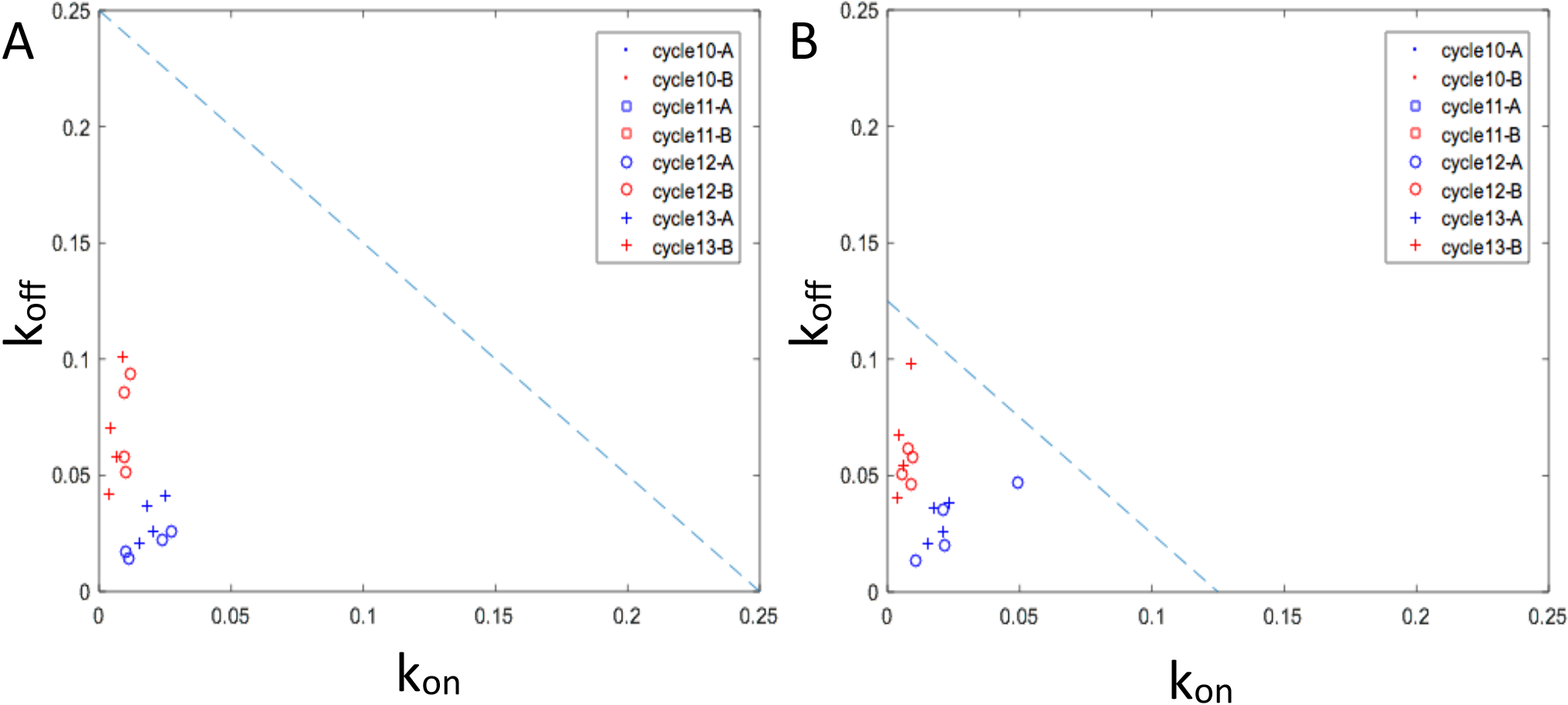
The dependence of the data fit on polymerase buffering time. Assuming different buffering times for the polymerase does not strongly affect the fit of the switching rates: a fit with τ_bufering_ = 4*s* (A) and τ_bufering_ = 8*s*. τ_bufering_ = 6*s* is used in the main text in Fig. 5D.

